# Paired plus-minus sequencing is an ultra-high throughput and accurate method for dual strand sequencing of DNA molecules

**DOI:** 10.1101/2025.08.11.669689

**Authors:** Alexandre Pellan Cheng, Itai Rusinek, Aaron Sossin, Adam J. Widman, Eti Meiri, Gat Krieger, Ori Hirschberg, Doron Shem Tov, Shlomit Gilad, Ariel Jaimovich, Omer Barad, Sammantha Avaylon, Srinivas Rajagopalan, Abigail G. Thress, Catherine Potenski, Tamara Prieto, Dennis J. Yuan, Rob Furatero, Alexi Runnels, Benjamin M. Costa, Jonathan E. Shoag, Majd Al Assaad, Michael Sigouros, Jyothi Manohar, Abigail King, David Wilkes, John Otilano, Murtaza S. Malbari, Olivier Elemento, Juan Miguel Mosquera, Nasser K. Altorki, Ashish Saxena, Margaret K. Callahan, Nicolas Robine, Soren Germer, Gilad D. Evrony, Bishoy M. Faltas, Dan-Avi Landau

## Abstract

Distinguishing real biological variation in the form of single-nucleotide variants (SNVs) from errors is a major challenge for genome sequencing technologies. This is particularly true in settings where SNVs are at low frequency such as cancer detection through liquid biopsy, or human somatic mosaicism. State-of-the-art molecular denoising approaches for DNA sequencing rely on duplex sequencing, where both strands of a single DNA molecule are sequenced to discern true variants from errors arising from single stranded DNA damage. However, such duplex approaches typically require massive over-sequencing to overcome low capture rates of duplex molecules. To address these challenges, we introduce paired plus-minus sequencing (ppmSeq) technology, in which both DNA strands are partitioned and clonally amplified on sequencing beads through emulsion PCR. In this reaction, both strands of a double-stranded DNA molecule contribute to a single sequencing read, allowing for a duplex yield that scales linearly with sequencing coverage across a wide range of inputs (1.8-98 ng). We benchmarked ppmSeq against current duplex sequencing technologies, demonstrating superior duplex recovery with ppmSeq, with a rate of 44%±5.5% (compared to ∼5-11% for leading duplex technologies). Using both genomic as well as cell-free DNA, we established error rates for ppmSeq, which had residual SNV detection error rates as low as 7.98×10^−8^ for gDNA (using an end-repair protocol with dideoxy nucleotides) and 3.5×10^−7^±7.5×10^−8^ for cell-free DNA. To test the capabilities of ppmSeq for error-corrected whole-genome sequencing (WGS) for clinical application, we assessed circulating tumor DNA (ctDNA) detection for disease monitoring in cancer patients. We demonstrated that ppmSeq enables powerful tumor-informed ctDNA detection at concentrations of 10^−4^ across most cancers, parts per million sensitivity in cancers with high mutation burden, and further increased sensitivity with higher sequencing depth. We then leveraged genome-wide trinucleotide mutation patterns characteristic of urothelial (APOBEC3-related and platinum exposure-related signatures) and lung (tobacco-exposure-related signatures) cancers to perform tumor-naive ctDNA detection, showing that ppmSeq can identify a disease-specific signal in plasma cell-free DNA without a matched tumor, and that this signal correlates with imaging-based disease metrics. Altogether, ppmSeq provides an error-corrected, cost-efficient and scalable approach for high-fidelity WGS that can be harnessed for challenging clinical applications and emerging frontiers in human somatic genetics where high accuracy is required for mutation identification.

## Introduction

Somatic SNVs have profound impact on human health and disease. Next generation sequencing (NGS) has transformed the discovery of SNVs across myriad clinical and research applications. However, the accuracy of current NGS approaches is constrained by pre-analytical and sequencing artifacts, mandating a reliance on multiple supporting fragments for a variant to be confidently called as a mutation. Therefore, to call SNVs with confidence, current NGS requires that a substantial proportion of cells harbor the variant, limiting the ability to identify variants with low allelic fraction represented by a single fragment. Such genome-scale applications are of emerging importance in both research exploration of the heterogenous somatic genome in non-malignant tissues^1–4^ and clinical interrogation of circulating cell-free DNA for cancer monitoring^5–7^.

Molecular barcoding and redundant sequencing have been harnessed to suppress error, lowering the number of unique molecules needed for variant calling. For example, duplex sequencing^8–11^, the current gold standard short-read error-correction strategy, requires that corresponding Watson & Crick strands be independently read and used to correct inconsistent base calls. These duplex sequencing protocols have been shown to have error rates between 10^−7^ and 10^−9^, employing various strategies to boost accuracy and duplex recovery^8–11^. For example, the bottleneck sequencing system^9^ (BotSeqS) and nanorate sequencing^8^ (NanoSeq) use a bottlenecking strategy, where input DNA is diluted up to 10,000 fold prior to DNA amplification to limit the number of unique double-stranded (ds)DNA molecules that undergo sequencing. Through bottlenecking, combined with extensive PCR and redundant sequencing, dsDNA recovery is improved over standard duplex protocols. However, extensive bottlenecking reduces the complexity of the library and limits the number of unique molecules that can be sequenced. To decrease error rates, NanoSeq additionally utilizes blunt-end restriction enzymes to fragment DNA or applies a single-strand exonuclease to remove single-stranded overhangs. The protocol also leverages dideoxy nucleotides (ddBTPs) to incorporate into nicked dsDNA and prevent amplification, eliminating the propagation of errors deriving from repair of abasic sites. Furthermore, Concatenating Original Duplex for Error Correction (CODEC)^11^ utilizes a novel quadruplex adapter and single primer extension to copy one strand of DNA onto the other, thereby creating two single-stranded molecules, each incorporating sequence information from both Watson and Crick strands, limiting the need to oversequence. However, the quadruplex adapter can bind to independent dsDNA molecules, reducing the final duplex yield. Currently, these approaches are limited to 5% (BotSeqS^9^, Nanoseq^8^, WGS duplex of cfDNA^6^) and 11% (CODEC^11^) read-to-duplex efficiency in whole-genome settings.

To circumvent this fundamental limitation of duplex sequencing, we introduce Paired Plus-Minus sequencing (ppmSeq), in which a PCR-free, high-throughput library preparation protocol leverages the hydrogen bonds between Watson & Crick strands to carry both DNA strands into a single clonal amplification reaction. Specifically, dsDNA is partitioned and clonally amplified through emulsion PCR on sequencing beads, which then undergo mostly-natural sequencing by synthesis. Thus, copies of the Watson strand and copies of the reverse-complement of the Crick strand are simultaneously read in a single sequencing read, allowing for high fidelity in SNV calls and a superior duplex recovery. Using ppmSeq for WGS of cell-free DNA, we demonstrate the ability to sequence low-input samples, showing that coverage scales linearly with input. We then present two key applications of this technology for rare variant discovery. First, we applied ppmSeq with enzyme blunting and ddBTPs to low mutational burden genomic DNA (gDNA) to establish our method’s error rate of 7.98×10^−8^ (7.78×10^−8^-8.18×10^−8^; 95% binomial confidence intervals, Wilson method^11^). Second, we applied ppmSeq to WGS of cell-free DNA to enable tumor-informed and tumor-naive cancer monitoring.

## Results

### ppmSeq encodes Watson and Crick strands into single sequencing reads and enables linearly scalable double-stranded DNA sequencing recovery

WGS is an exciting strategy for genome-wide variant identification, but high-throughput sequencing technologies are too error prone (∼1 error in 1,000 bases without analytical denoising and ∼1 error in 10,000 bases with denoising^6^) for rare variant identification (i.e., low variant allele frequency (VAF) variants supported by 1-2 sequencing reads in a sample). Moreover, in many instances, errors arise not only in sequencing but also due to DNA degradation or nicking, often resulting in error rates limiting accuracy of NGS methods based on reading one DNA strand (∼10^−5^ errors per base pair)^8^. While duplex sequencing is the gold standard error correction strategy for short reads^8–11^, the likelihood of sequencing both strands of the same dsDNA molecule in whole genome settings is low. Though dsDNA yield can be improved to ∼5% with bottlenecking strategies^8^, these recovery rates substantially limit sensitivity.

To improve the recovery of dsDNA molecules, we sought a technology that could encode dsDNA information into single sequencing reads. The Ultima Genomics platform uses an emulsion PCR clonal amplification step that does not require prior DNA denaturation^12^ and is able to encode single duplexes into individual sequencing reads. The key principle is that mismatched bases on a given dsDNA molecule would cause a conflicting sequencing signal at a given position and result in a low base quality at that position, which can be filtered bioinformatically. To quantify the ratio of Watson copies and Crick copies on the sequencing bead, a critical innovation involves custom adapter sequences containing a known mismatch sequence (here, a 6-bp sequence of adenines on both strands of the adapter). Given the mismatch, the adapter sequence of Watson copies is different than the one on Crick copies, resulting in a pattern on the adapter sequence that can be used to quantify the ratio between copies of both strands on the sequencing bead (**Figure 1A, Supplementary Figure 1**).

**Figure 1.**
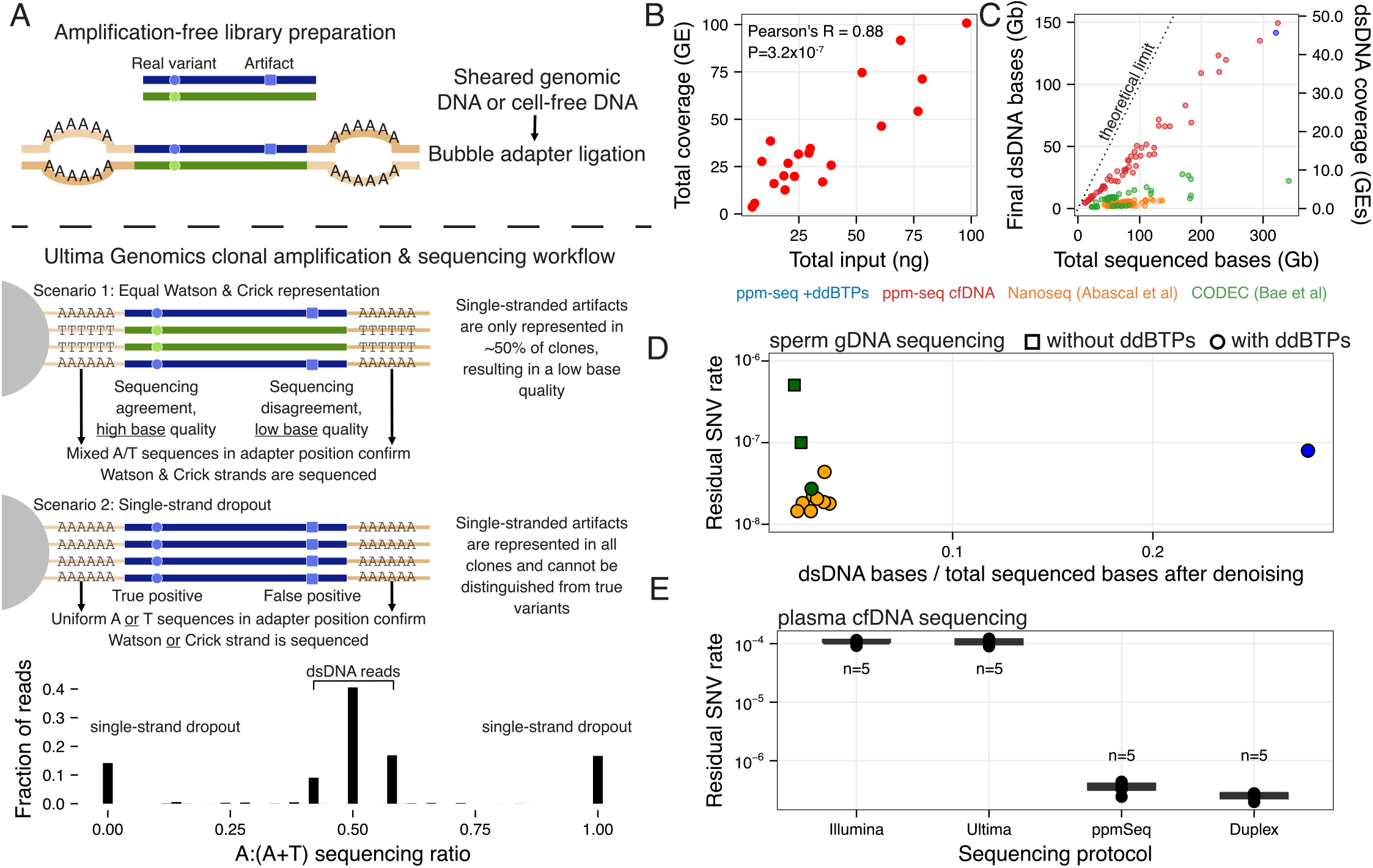
Paired plus-minus sequencing (ppmSeq) enables high fidelity duplex sequencing and linearly scalable double-stranded (ds)DNA sequencing recovery. **A**. ppmSeq workflow. Genomic DNA (gDNA; sheared) or cell-free DNA (cfDNA; unsheared) undergoes end-repair and adapter ligation. The adapter contains a known sequence of 6 non-complementary base pairs (here, As on each strand). Following library preparation, adapter-ligated molecules undergo clonal amplification through emulsion PCR, creating distinguishable Watson and Crick strands via the different adapter sequences. **Scenario 1: Equal representation of Watson & Crick strands**. During adapter sequencing, the mismatched sequences allow positive identification of dsDNA (termed mixed) reads. Damaged bases (illustrated as squares) can cause polymerases to incorporate the wrong base during emulsion PCR, leading to mismatched sequences during template sequencing. The conflicting sequencing signal at the mismatch position results in the encoding of that base with a low quality. True variants (illustrated as circles) are amplified from both native strands and do not cause conflicting sequencing signal, resulting in a high base quality encoding. **Scenario 2: Single-strand dropout**. Damaged bases (illustrated as squares) are amplified throughout the single-stranded clones and are indistinguishable from true variants (illustrated as circles). Here, both true variants and false positives are encoded with high base quality scores and cannot be readily distinguished. However, the sequencing signal at the adapter positions will also be consistent (uniform A or T reads), allowing for the identification of these reads and their computational filtering. **Bottom**: Sequencing signal at the adapter positions for cell-free DNA sample UPN-035. Single-stranded reads can be identified through the consistent A or T sequencing signal, and dsDNA is identified through mixed A:T signal. **B**. Assessment of total genome coverage [genome equivalents (GE)] across different total input amounts (ng) of cell-free DNA (cfDNA) sequenced using ppmSeq. There is a high correlation between input and coverage. **C**. Comparison of mixed read yields (representing duplex reads) obtained from ppmSeq libraries (unsheared cell-free DNA or sheared genomic DNA) with reported duplex yields from published unique molecular identifier technologies (NanoSeq and CODEC). Chain-terminating dideoxy bases (ddBTPs; used in NanoSeq) are incorporated into nicked dsDNA to prevent amplification and improve duplex error rates for the gDNA ppmSeq libraries. Duplex recovery is assessed by the final number of dsDNA bases sequenced (in Gb) and total dsDNA coverage (in genome equivalents) across total bases sequenced. ppmSeq provides the highest yield of dsDNA recovery compared to available methods for cell-free DNA and gDNA (with ddBTPs). **D**. Residual SNV rate from ppmSeq libraries (mixed reads only) compared to SNV rate from published duplex sequencing protocols for sperm gDNA libraries, showing the fraction of duplex reads recovered for each technology. For ppmSeq libraries, the gDNA sample was sequenced to 100x, and variants that occurred in a single read were considered errors. Squares represent samples that were not prepared with ddBTPs during end-repair, thus enabling the amplification of nicked and overhanging DNA. Circles represent samples that were prepared with ddBTPs. **E**. Residual SNV rates of cell-free DNA libraries (n = 5 for each method, mixed reads only) generated with standard whole-genome sequencing (Illumina NovaSeq 6000 and Ultima UG 100), ppmSeq (Ultima) and duplex (Ultima, Cheng et al. *Nature Methods*, 2025^6^). SNV rates were measured on the scale of the entire genome, where any single occurring variant is considered an error. Samples were not matched for each method, and sequencing statistics are available in **Supplementary Table 1**.

To establish ppmSeq as an approach for accurate detection of low-frequency variants, we first assessed scalability in the context of WGS of cell-free DNA, which has been shown to be a promising biomarker for infection detection^13–16^, non-invasive prenatal testing^17,18^, ultrasensitive cancer monitoring^5,7,19–25^, and other clinical applications. We reasoned that because molecules of interest in cell-free DNA are often present in extremely low abundance, and cell-free DNA is scarce (roughly 1,000-10,000 genomic equivalents [GEs] per milliliter of plasma^26^), the combination of high accuracy and efficiency enabled by ppmSeq would make it a uniquely effective tool for variant calling in this context.

To demonstrate the scalability of ppmSeq in terms of DNA input versus sequencing coverage output, we performed exhaustive sequencing of cell-free DNA with various input amounts (range 1.8-98.0 ng, PCR free) and measured the total coverage obtained (**Figure 1B, Supplementary Figure 2A**). We tested the ppmSeq chemistry prototype on three cohorts (cancer-free controls, patients with lung cancer and patients with urothelial cancer, see **Methods** and **Supplementary Table 1**), and obtained average total coverage to input ratios of 1.3±0.7, 1.2±0.7 and 2.2±0.7 for the cancer-free control, lung cancer and urothelial cancer cohorts, respectively (±standard deviation, in units of GEs obtained per ng of cell-free DNA). The commercially available ppmSeq kit (UG ppmSeq™ Cell-Free DNA Solaris Free ppmSeq D1000986 Rev.02) demonstrated a total coverage to input ratio of 22.2±2.9 (**Supplementary Figure 2B**). Thus, ppmSeq enables low-input sequencing where total coverage scales linearly with input amounts.

### ppmSeq reads maintain fidelity approximating duplex sequencing with an order of magnitude greater yield

To benchmark ppmSeq performance, we compared our protocol’s dsDNA molecule recovery with that of previously published, state-of-the-art short-read sequencing protocols, NanoSeq^8^ and CODEC^11^. We first assessed dsDNA recovery from cell-free DNA, finding that ppmSeq provides the highest yield of dsDNA recovery, with a rate of 44%±5.5% in cell-free DNA, which was significantly higher than the published results from application of CODEC and the application of NanoSeq to gDNA (11%±5.7% and 5%±1.4%, respectively; **Figure 1C, Methods**). ppmSeq dsDNA yield was positively correlated with total sequencing yield (Pearson’s R = 0.98, *P* < 2.2×10^−16^).

Next, to evaluate the fidelity of ppmSeq reads and determine error rates, we prepared a custom library of low mutational burden sample, using ddBTP blocking to decrease the effect of pre-sequencing artifacts, as in the NanoSeq protocol. Specifically, sperm gDNA was fragmented with the HpyCH4V restriction enzyme to provide blunt ends gDNA, and ddBTPs were added to inhibit the amplification of nick-containing dsDNA. We reasoned that utilizing sperm gDNA, which has the lowest mutational burden of any human tissue, in conjunction with error-resistant library preparation techniques^8,27^, would enable an accurate error rate estimation of the ppmSeq platform. The gDNA sample was sequenced to ∼100x and total dsDNA recovery was 44%. After quality recalibration and analytical denoising (**Methods, Supplementary Figures 3**,**4**,**5, Supplementary Tables 2**,**3**), we achieved an error rate of 7.98×10^−8^ and a dsDNA recovery of 28% (**Figure 1D**). The ddBTP-blocked ppmSeq library error rate (7.98×10^−8^) was higher than error rates from NanoSeq (2.13×10^−8^±9.5×10^−9^, all ddBTP blocked, n = 8) and CODEC libraries (n = 3; 2.72×10^−8^ with ddBTP blocking and 1.0×10^−7^, 5.07×10^−7^ without), but with a much higher denoised yield (28% dsDNA bases / total bases sequenced for ppmSeq versus 3.1%±0.6% and 2.5% ±0.4% post-denoising for NanoSeq and CODEC libraries, respectively; **Figure 1D, Methods**).

Encouraged by the high fidelity reads afforded by the ppmSeq platform, we next sought to estimate the effective error rate of ppmSeq in cell-free DNA. In the cell-free DNA of healthy individuals, mutations can arise as a function of biological phenomena (for example, SNVs affecting normal blood cells), artifacts from library preparation or sequencing, or contamination. Therefore, in cancer detection studies, cell-free DNA from cancer-free individuals is often used as a control from which background rates can be established. As cell-free DNA is highly degraded and short (∼167 bp)^28^, with most dsDNA molecules containing overhangs, it is not amenable to the strategies for error suppression used in the NanoSeq protocols. For example, restriction enzyme digestion, which would require two digestion sites per cell-free DNA molecule, cannot be used for blunting the short fragment cell-free DNA. In addition, application of mung bean exonuclease^8^ to create blunt ends would be too degradative for low input cell-free DNA applications. Further, the use of ddBTPs would inhibit the amplification of overhang-containing dsDNA, which would limit the sequencing of most cell-free DNA molecules. Therefore, we performed standard, PCR-free library preparation of cell-free DNA, followed by ppmSeq and measured residual SNV rates on the scale of the entire genome, where any single occurring variant was considered to be an error. We compared ppmSeq error rates in cell-free DNA to standard Illumina, standard Ultima or duplex sequencing with Ultima^6^. Standard Illumina and Ultima libraries showed the highest residual SNV rates (1.07×10^−4^±9.29×10^−6^ and 1.06×10^−4^±1.30×10^−5^, respectively). ppmSeq exhibited significantly lower error rates (3.5×10^−7^±7.5×10^−8^ [mean ± standard deviation]), on par with duplex sequencing (2.5×10^−7^±3.09×10^−8^) (**Figure 1E**), and comparable to the CODEC error rate in application to cell-free DNA (2.9×10^™7^)^11^.

### Application of ppmSeq to tumor-informed ctDNA detection

ctDNA has emerged as a promising biomarker for the non-invasive detection and monitoring of cancer. Our previous work^5^ has shown that high-throughput targeted sequencing rapidly exhausts available genomes for sequencing (1,000-10,000 GEs per mL of plasma), which sets a hard ceiling on ctDNA detection. In contrast, WGS approaches can exploit breadth of coverage to supplant depth, eliminating the reliance on the detection of few sites, to enhance ctDNA characterization in low tumor fraction settings.

Given the low error rate of ppmSeq when used for WGS of cell-free DNA, we next sought to establish ppmSeq for ctDNA detection through plasma WGS, reasoning that this approach would enable accurate detection of low tumor fractions. To first benchmark tumor fraction detection with ppmSeq, we profiled an *in vitro* mixture of Genome in a Bottle (GIAB^29^) cell lines (**Figure 2A**), where HG001 and HG005 were both diluted into HG002 DNA in proportions of 1, 10 and 100 parts-per-million (n = 3 replicates). By measuring the unique HG001 and HG005 germline mutation (homozygous single-nucleotide polymorphisms (SNPs) were subsampled to 30K to model cancer SNVs) presence in the diluted samples, we observed high concordance of measured and expected GIAB fractions with low background rate (**Figure 2A, Supplementary Figure 6** and **Supplementary Tables 4, 5**), indicating that ppmSeq would be able to detect tumor fractions in the parts-per-million range.

**Figure 2.**
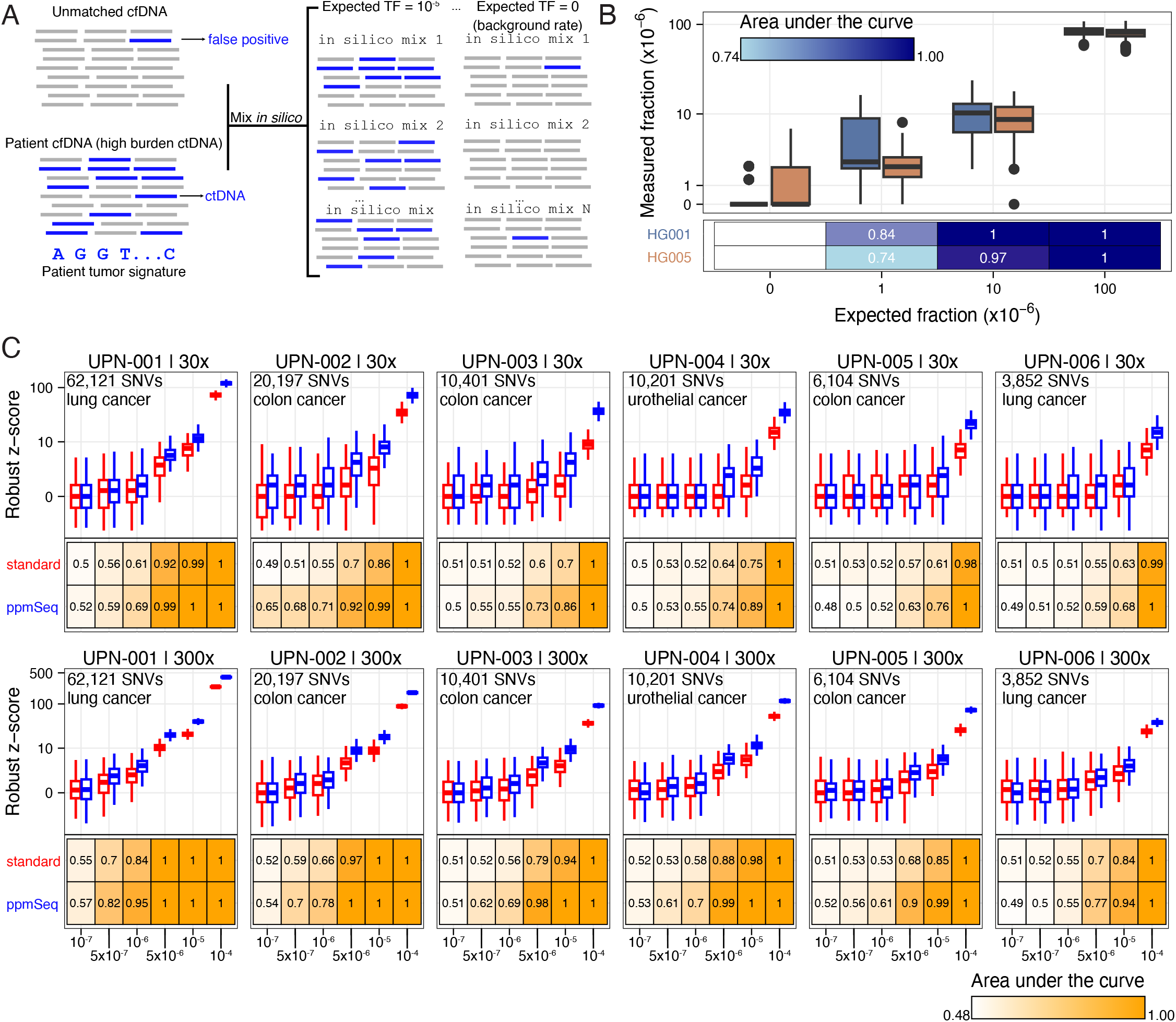
*in silico* mixing studies for tumor-informed ctDNA detection demonstrates superior performance of ppmSeq compared to standard whole-genome sequencing on a UG 100 sequencer. **A**. *in silico* mixing workflow. Briefly, patient cfDNA is computationally mixed into unmatched cfDNA from cancer-free controls in known proportions to generate a range of expected tumor fractions (TF). Clinical patient information, cancer types and tumor fractions are available in **Supplementary Table 6** and **7**. **B**. Benchmarking tumor fraction detection with ppmSeq using an in-vitro mixture of Genome in a Bottle (GIAB^29^) genomic DNA. Each of the HG001- and HG005-unique homozygous variants were subsampled for a signature of size 30,000 SNVs, for 30 times with repeats. The distribution of GIAB-supporting reads is shown in boxplots, the area under ROC-curve, for each sample against the HG002 sample (expected fraction=0) is shown in heatmaps. Results per replicate are presented in **Supplementary Figure 2B**. **C**. *in silico* mixing study comparing matched libraries that underwent both standard Ultima sequencing (amplified library using KAPA HYPER PREP kit (KK8505)) and Ultima ppmSeq from a cohort consisting of cancer patients (n = 6) with matched plasma, tumor and normal tissue sequencing at 30x (top row) and 300x (bottom row) total coverage. To generate cancer-specific mutation profiles for each patient, tumor and normal samples were sequenced [average of 100x (72-132x) and 80x (64-146x), respectively]. Plasma samples were sequenced to an average of 70x (28-103x) for ppmSeq and 139x (110-166x) for standard Ultima. Patient-specific mutational profiles were computationally mixed into an unmatched cfDNA profile (i.e., cell-free DNA from another patient) and reads (mixed and non-mixed) matching the mutational profile were tabulated. For each simulated condition, we drew 1,000 samples and computed a robust z-score as the difference between the matched and background distributions. The average results across controls yielded a calibrated measure of ctDNA detectability, informing the threshold at which true tumor-derived signals can be confidently distinguished from artifacts. Area under the receiving operator curve (AUC) values demonstrates detection performance at the different admixed TFs versus negative controls (TF = 0) as measured by z score. Mixed and non-mixed reads were used for this analysis.

Matched tumor mutational profiles can inform genome-wide tumor SNV detection in cell-free DNA, such that, unlike targeted panels, the available number of GEs is no longer the limiting factor for successful ctDNA detection. To test performance of ppmSeq for ctDNA detection through tumor-informed WGS, we evaluated ctDNA detection and background error rates using *in silico* dilution experiments, as we had performed in our previous evaluation for ctDNA detection^7^. Briefly, previous evaluation showed that Illumina and Ultima sequencing platforms (Illumina NovaSeq 6000 and Ultima UG 100) performed similarly in terms of ctDNA detection and background error rates by sequencing a patient with readily detectable ctDNA, and performing an *in silico* dilution with sequencing reads from unmatched cell-free DNA (i.e., from another individual). To assess ppmSeq, we performed a similar experiment (**Figure 2B**), comparing matched libraries that underwent both standard amplified Ultima sequencing and Ultima ppmSeq. Our cohort consisted of six lung, bladder and colorectal cancer patients (unique patient number [UPN]-001-UPN-006, **Supplementary Table 6**) with matched plasma, tumor and normal tissue sequencing.

Tumor and normal samples were sequenced to an average of 100x (72-132x) and 80x (64-146x), respectively, and ppmSeq and standard plasma samples were sequenced to an average of 70x (28-103x) and 139x (110-166x), respectively. Tumors harbored between 3,852 and 62,121 SNVs, representing a range of mutational burden observed in solid cancer from average (∼1/Mb) to high (>10/Mb)^30^. Plasma tumor fractions were estimated by counting the number of cell-free DNA fragments carrying a tumor-matching mutation and dividing by the total number of reads overlapping with a tumor-mutated locus, as previously described^5–7^. Estimates from standard and ppmSeq sequencing showed strong agreement (R^2^ = 0.998, Pearson’s R = 0.9999, *P* = 1.72×10^−6^, **Supplementary Figure 7, Supplementary Table 7**) and ranged from 0.001-6%.

To compare low-burden ctDNA detection in standard sequencing and ppmSeq, we performed *in silico* mixing studies by diluting ctDNA datasets with unmatched cell-free DNA at known ratios^5^ (**Methods, Figure 2B**). We generated admixtures for each of the 6 patients (n=1 urothelial cancer; n=2 lung cancer; n = 3 colon cancer; **Supplementary Table 8** and **6, Figure 2C**) at tumor fractions ranging from 10^−7^ to 10^−4^ (n = 1,000 replicates per patient, per tumor fraction), and assay performance was evaluated by receiving operating curve analysis by comparing the ctDNA detection rate of replicates at a given tumor fraction to the ctDNA detection rate of cross-patient controls. First, we analyzed samples at 30x total sequencing depth. Detection at 10^−4^ was achieved with AUC = 1 across all cancer samples using ppmSeq, including the one with the lowest mutation burden (3.8K), consistent with the limit of detection of commercially-available tumor-informed panel assays^31^. Of note, ∼70% of solid tumors in adults have a mutation burden higher than 4K^32^. ctDNA was more readily detectable in concentrations of 10^−5^ in ppmSeq datasets from patients with 10,000 or more SNVs (corresponding to ∼35% of solid tumors^32^), compared to standard sequencing (mean AUC of 0.94 (range 0.86-1.00) for ppmSeq datasets and 0.83 (range 0.77-0.99) for standard sequencing datasets, **Figure 2C, Supplementary Table 8**).

Encouraged by these results at 30x sequencing coverage, we sought to evaluate whether deeper sequencing coverage would enable lower limits of ctDNA detection, as ctDNA has been shown to be at levels at or below 10^−5^ in early-stage and minimal residual disease scenarios^25,33^ and might not be sampled at 30x sequencing coverage^5,6^. Therefore, we reasoned that ultra-deep, highly accurate sequencing would be required to detect ultra-low ctDNA burden and explored the potential impact of 300x WGS in improving ctDNA sensitivity. To simulate performance at this coverage, we used a parametric bootstrapping approach, where the noise rate was evaluated based on the lower coverage data and reads were sampled using a Poisson distribution with parameters corresponding to the higher coverage. As expected, increased sequencing depth resulted in improved ctDNA detection, with robust detection of 10^−5^ ctDNA concentration across all samples (**Figure 2C**). Excitingly, for the highest mutation burden tumor (62K, as is commonly observed in malignancies such as melanoma, lung and mismatch repair deficient cancers), ctDNA was even detectable in ppmSeq datasets at ctDNA concentrations as low as 5×10^−7^ (AUC = 0.82), suggesting that deep, error-corrected WGS could enable the detection of MRD in high mutational burden tumors at ctDNA levels far below the sensitivity floor of current clinical assays^31^. This is particularly relevant given that ctDNA positivity after curative-intent treatment is strongly associated with disease recurrence^22^. Notably, improving and ctDNA assay sensitivity from 10^−4^ to 10^−6^ has been shown to recover additional clinically high-risk patients missed by prior assays^25^, with prognostic value^34^. Together, these results highlight the ability of ppmSeq to enhance tumor-informed ultra-low ctDNA detection in patients with cancer.

### ppmSeq empowers tumor-naive (plasma-only) ctDNA detection

Encouraged by the high scalability of ppmSeq and its low background error rate, we sought to investigate its applicability to tumor-naive ctDNA detection. This is of significant clinical importance, as access to and scarcity of matched tumor tissue limits tumor-informed cancer monitoring^35–37^. We have previously reported that whole-genome duplex sequencing of cell-free DNA was a potential tool for detecting ctDNA without matched tumors, but at the cost of massive over-sequencing^6^. Specifically, we have shown that duplex WGS could be used to deconvolute mutational signatures arising from multiple biological processes, such as those originating from ultraviolet exposure mutagenesis in patients with melanoma or APOBEC3-derived processes in patients with urothelial cancer, and that the presence or absence of these signatures could identify and quantify the degree of disease. Given the similar error rates between ppmSeq and duplex WGS, we reasoned that ppmSeq could be used for mutational signature-based ctDNA detection, but without the over-sequencing that is required for traditional duplex WGS. To test this, we generated ppmSeq datasets from 20 cell-free DNA samples from 20 patients with urothelial cancer (stages II-IV) for whom duplex WGS was generated as part of previously published work^6^. Of these 20 patients, 11 had previously received neoadjuvant platinum chemotherapy, and 13 had tumor tissue available for sequencing (**Figure 3A, Supplementary Table 9**).

**Figure 3:**
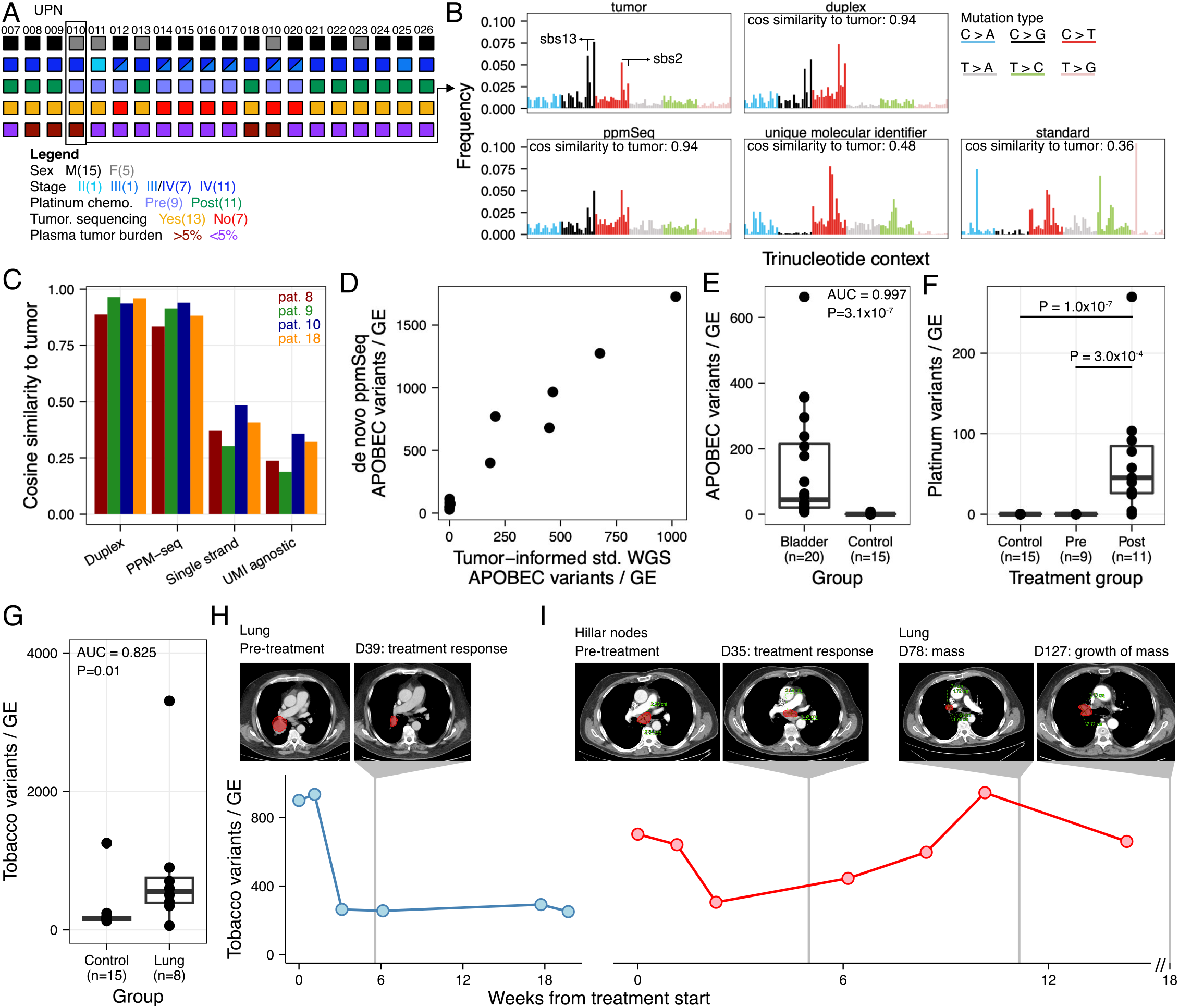
ppmSeq empowers tumor-naive ctDNA detection. **A**. Cohort description for tumor-naïve ctDNA detection comprising 20 patients with urothelial cancer (stages II-IV). 20 cfDNA samples were sequenced with ppmSeq and matched Ultima duplex whole-genome sequencing. Of these 20 patients, 11 had previously received neoadjuvant platinum chemotherapy, and 13 had tumor tissue available for sequencing datasets. **B**. Comparison of the trinucleotide frequencies obtained from matched duplex whole-genome sequencing with different levels of denoising (duplex: duplex consensus variant calling; unique molecular identifier: strand-agnostic unique molecular identifier (UMI) consensus variant calling; or standard: UMI-agnostic denoising) and ppmSeq (mixed reads only). Trinucleotide frequencies of each variant-calling method were compared to those of the matched tumor mutational pattern using cosine similarities. SBS2 and SBS13 are prominent in urothelial cancers and represent APOBEC3-associated signatures reflecting cytidine deaminase activity, where C>T transitions can occur as a function of uracil generation by cytidine deaminase activity (SBS2) and well as C>G and C>A mutations that can arise as a result of polymerase errors following uracil excision (SBS13)^40^. **C**. Cosine similarities of trinucleotide frequencies in cfDNA compared to those from matched tumor across sequencing strategies (duplex: duplex consensus variant calling; single strand: strand-agnostic unique molecular identifier (UMI) consensus variant calling; UMI-agnostic denoising) and ppmSeq (mixed reads only) for 4 patients with high tumor burden (defined as having a tumor fraction >5% as measured through ichorCNA) and matched tumor sequencing. **D**. Comparison of tumor-naive APOBEC3 scores generated from cfDNA (de novo ppmSeq; mixed reads only) versus tumor-informed APOBEC3 scores (standard whole-genome sequencing of tumor in the n=13 patients for whom tumor sequencing was available. Tumor-informed ctDNA based tumor fractions were measured by counting variants found in the ctDNA that matched the tumor sequencing profile, and by dividing by the total number of reads overlapping tumor-specific mutations. The tumor-informed APOBEC3 score is obtained by multiplying the tumor fraction by the relative contribution of APOBEC3 mutations (SBS2 + SBS13) measured from the tumor mutational profile. There is a strong correlation between the expected, tumor-informed APOBEC3 contributions in the plasma and the de novo APOBEC3 score. The de novo APOBEC3 score is obtained by fitting the trinucleotide profiles of somatic variants to a curated catalog of cancer-associated SBS signatures (see **Methods**). **E**. Plasma mutational scores from APOBEC3 variants (SBS2 + SBS13) in individuals with urothelial cancer (stage II–IV; n = 20) and healthy individuals (n = 15). Plasma signatures were fit to a custom reference of signatures comprising SBS2, SBS13, SBS31, SBS35 (Cosmic v3.3), clonal hematopoiesis^41^ and one derived from a subset of cancer-free controls (**Methods**). For E-I, only mixed reads were used to measure plasma mutational scores. **F**. Plasma mutational scores from platinum therapy mutagenesis (SBS31) in individuals with urothelial cancer (stage II–IV; n = 11 after, n = 9 before) and healthy individuals (n = 15). Plasma signatures were fit to a custom reference of signatures comprising SBS2, SBS13, SBS31 and SBS35 (Cosmic v3.3), clonal hematopoiesis^41^ and one derived from a subset of cancer-free controls (**Methods**). **G**. Plasma mutational scores from tobacco variants (SBS4) in individuals with lung cancer (n = 8) and healthy individuals (n = 15). Plasma signatures were fit to a custom reference of signatures comprising SBS4 (Cosmic v3.3), clonal hematopoiesis^41^ and one derived from a subset of cancer-free controls (**Methods**). **H**. Serial plasma monitoring of patient UPN-028 with ppmSeq and no matched tumor corresponds to changes seen on imaging. Pre-treatment and post-treatment (day 39) CT imaging of a lung cancer patient with decreased ctDNA (tobacco signature obtained from cfDNA sequencing with ppmSeq) in response to immune checkpoint inhibition therapy. **I**. Serial plasma monitoring of patient UPN-030 with ppmSeq and no matched tumor corresponds to changes seen on imaging. Pre-treatment and post-treatment (day 35) CT imaging of Hillar nodes and day 78 and day 127 CT imaging of lung a lung cancer patient reflects ctDNA levels (tobacco signature obtained from cfDNA sequencing with ppmSeq) in response to immune checkpoint inhibition therapy.

First, to benchmark ppmSeq ability for mutation calling based on a single-supporting read, we compared matched unique molecular identifier (UMI)-based duplex WGS and ppmSeq cell-free DNA libraries for four patients with readily detectable ctDNA, as defined by a tumor fraction greater than 5% determined by ichorCNA^38^, with tumor and normal tissue also available to generate ground truth mutational profiles. In duplex libraries, somatic mutations were called on single molecules using either *i)* duplex consensus variant calling; *ii)* strand-agnostic UMI consensus variant calling or *iii)* standard (UMI-agnostic) denoising as previously described^6^. ppmSeq somatic mutations were identified using recalibrated variant qualities (**Methods**). We compared the trinucleotide frequencies of each variant-calling method to that of the tumor mutational pattern using cosine similarities (**Figure 3B;** patient UPN-010). We found that the mutational profiles of ppmSeq and duplex are highly similar to the somatic mutations found in the matched tumor biopsies (0.89±0.05 and 0.94±0.04 cosine similarities, respectively), and significantly higher than that of other denoising methods (0.39±0.08 and 0.28±0.08, for strand-agnostic UMI and standard analytical denoising, respectively) across four patients (**Figure 3C**). In addition, the trinucleotide mutational profiles of the tumor, duplex and ppmSeq datasets could be readily mapped to COSMIC signatures, predominantly reflecting APOBEC3-induced mutagenesis and platinum chemotherapy exposure, consistent with the known mutational processes of urothelial cancer^39^ and treatment histories of the patients (**Figure 3A, Supplementary Figure 8**).

We then tested the ability of ppmSeq to identify mutational signatures to detect ctDNA without the need for tumor sequencing, as previously described^6^. Briefly, we tabulated the trinucleotide profiles of cell-free DNA somatic variants and fit the observed mutational profile to a curated reference of known mutational profiles (**Methods, Supplementary Table 9**). To evaluate our tumor-naive method’s ability to accurately quantify ctDNA, we compared our tumor-naive scores to tumor-informed tumor fractions in the n = 13 patients for which tumor sequencing was available. Specifically, tumor-informed ctDNA based tumor fractions were measured by counting variants found in the ctDNA that matched the tumor sequencing profile, and by dividing by the total number of reads overlapping tumor-identified mutation loci. Tumor-informed APOBEC3 contributions were then obtained by multiplying the tumor fraction by the relative contribution of APOBEC3 mutations to the tumor’s mutational profile, as previously described^6^. For tumor-naive detection, germline variants were removed using the normal sequencing datasets. We found a strong correlation between the expected, tumor-informed APOBEC3 contributions in the plasma and our de novo APOBEC3 score (Pearson’s R = 0.98, *P* = 8.12×10^−9^, **Figure 3D**). To extend the analysis to samples without matched normal sequencing, we retabulated de novo APOBEC3 scores for all samples using a 0.2 allele frequency cutoff to remove germline variants. These de novo APOBEC3 score readily separated patients with urothelial cancer from cancer-free controls (AUC = 0.997, *P* = 3.1×10^−7^, Wilcoxon test, **Figure 3E, Supplementary Table 10**), despite the low tumor burden in some plasma samples (n = 7 below tumor fractions of 0.001; n = 6 above 0.005 for the 13 patients with tumors available for tumor-informed tumor fraction estimations^6^). Moreover, we identified a significant increase in the contribution of platinum chemotherapy-induced mutations (SBS31 contributions) to the trinucleotide profiles of patients who received chemotherapy prior to sampling, but not in controls or chemotherapy unexposed patients (**Figure 3F**, mean of 67.4±75.0 platinum variants / GE in chemotherapy-treated patients and no detections in pre-treatment and cancer-free controls; *P* = 1.0×10^−7^ between chemotherapy-treated patients and cancer-free controls and *P* = 3.0×10^−4^ between treated and chemotherapy-naive patients).

Finally, we tested our tumor-naive ppmSeq ctDNA detection for the monitoring of lung cancer patients receiving neoadjuvant treatment. In this clinical scenario, tumors have not yet been resected, limiting the applicability of tumor-informed ctDNA detection based on small biopsies. We reasoned that ctDNA could be detected through the presence of tobacco-induced DNA mutagenesis (SBS4). Here, SBS4-derived variants were significantly enriched in pre-treatment lung cancer patients (n = 8; all smokers or former smokers (n = 5 non-small cell lung cancer and n = 3 small-cell lung cancer. **Supplementary Table 8**) compared to cancer-free controls (n = 7 smokers; n = 5 non-smokers; n = 3 unknown, **Figure 3G**, mean of 853±1023 and 237±283 tobacco-derived variants per GE, respectively, AUC = 0.83, *P* = 0.01, Wilcoxon, **Supplementary Table 11**), demonstrating the feasibility of utilizing tobacco-derived mutations to identify ctDNA from lung cancer patients. Interestingly, within the cancer-free control group, smokers had an elevated number of tobacco-derived variants than non-smokers (334±407 versus 158±12 tobacco-derived variants per GE), but this difference was not significant (*P* = 0.11, Wilcoxon). To explore the possibility of tumor-naive (plasma-only without matched tumors) monitoring in these patients, we applied ppmSeq to longitudinal plasma samples from two small-cell lung cancer patients receiving immune checkpoint inhibitors (**Figure 3H**,**I**). ctDNA dynamics correlated with and preceded imaging modalities, reflecting initial treatment response. In patient UPN-028 (**Figure 3H**), an initial response to treatment can be observed as measured through ctDNA and a decrease in tumor volume on imaging. In patient UPN-030, a similar response to treatment can be observed in the ctDNA and imaging on the hilar nodes. However, ctDNA increased consistently 6 weeks after treatment start, and growing tumor mass could be observed in the lungs. These patient vignettes highlight the ability of ppmSeq to monitor patients without the requirement for tumor sequencing.

## Discussion

A perennial challenge in genomics science is developing DNA sequencing approaches that accurately identify true SNVs, with experimental or analytical strategies that are capable of distinguishing real signals from errors. The next-generation sequencing revolution has empowered dramatic expansion of genomics applications at unprecedented scale and breadth, with continuous improvements in sensitivity and accuracy pushing the boundaries of detection to even ultra-low frequency variants. These capabilities have advanced biological discoveries about mutation landscapes in humans^42,43^, have been harnessed to define off-target effects of genome editing enzymes^44^, and have provided clinical insights into human disease, including the identification of subclonal mutations in cancer^45–47^ and tracking disease burden through ctDNA analysis^5–7,20,25,48–53^. As our understanding of the somatic genome evolves, the ability to accurately call ultra-low frequency variants has important applications across human health and disease, with the potential to transform our understanding of cancer, chronic disease and aging^1,6,54–57^.

Towards this aim, recent strategies for boosting sequencing accuracy to confidently call ultra-low frequency SNVs involve duplex sequencing, where both the top and bottom strands of DNA are sequenced independently, leveraging the intrinsic complementary nature of the molecule to distinguish true SNVs from single-strand errors. While this approach increases specificity of detection of ultra-rare variants, there are tradeoffs in coverage and cost. For example, multiplexed PCR-based strategies (including Pro-Seq^58^ and SaferSeqS^19^) employ deep sequencing of targeted panels, allowing for highly sensitive detection but only of a limited number of loci. Sampling dilution strategies such as those used in BotSeqS^9^ and Nanoseq^8^ increase recovery of duplex molecules, but this bottlenecking reduces sensitivity. NanoSeq provides extremely low error rates (1 in 10^−9^) through use of bottlenecking, blunt-end restriction enzymes and dideoxy nucleotides for repair, allowing for highly specific sequencing of ∼30% of the genome^8^, but is not suitable for profiling highly fragmented templates such as cell-free DNA, where using ddBTPs to block the amplification of sticky-ended molecules would result in extremely low yields. Adaption of this NanoSeq using blunt-end nucleases expands the genomic regions that can be profiled in higher input samples, and more recently an exome-wide version has been developed^59^. However, despite these advances, broad implementation of duplex whole-genome sequencing has been inhibited by high cost, given the need for oversequencing to reliably capture duplex molecules (up to 100-fold excess reads^11^).

To address this challenge, we leveraged ppmSeq, which delivers the advantage of scalable, high-accuracy dsDNA sequencing at low cost, through increasing dsDNA yield and eliminating the requirement for high levels of oversequencing. This is accomplished through flow-based sequencing and by using custom mismatching adaptors to encode single duplexes into individual sequencing reads, for high coverage whole-genome sequencing from low-input samples. Here, dsDNA molecules are confidently identified by using 6-nt mismatching adaptors and base calling quality metrics to definitively assess Watson and Crick strand representation. The PCR-free, high-throughput library preparation workflow takes advantage of the hydrogen bonds between dsDNA strands to carry both strands into a single clonal amplification reaction. Subsequent clonal amplification and sequencing allows each strand of a dsDNA molecule to be encoded as a single sequencing read. We benchmarked ppmSeq against state-of-the-art short-read sequencing methods, demonstrating superior dsDNA strand recovery using ppmSeq (44±5.5%) compared with NanoSeq (5±1.4%) or CODEC (11±5.7%) (measured prior to denoising). Using ddBTPs to inhibit the amplification of nick-containing dsDNA together with a read-centric denoising classifier, we applied ppmSeq to sperm gDNA and achieved an error rate of 7.98.x10^−8^. This rate is comparable, albeit somewhat higher than NanoSeq and CODEC (∼2 × 10^−8^), yet dsDNA strand recovery after analytical denoising remains an order of magnitude higher (28% vs. 3.1% and 2.5% for NanoSeq and CODEC, respectively). Thus, ppmSeq provides high dsDNA recovery together with low error rate for efficient high-accuracy whole-genome sequencing that can analyze low-input samples.

Given these abilities, we reasoned that ppmSeq would be particularly well suited to application in ctDNA detection from plasma whole-genome sequencing, where signal is sparse and it is highly challenging to distinguish true variants from error. In such cases, the whole-genome sequencing approach is advantageous over deep sequencing of targeted panels^5,7^. Rather than amplification of specific loci that can be missed or rapidly exhausted from limiting amounts of template, aggregate genome-wide signal can be assessed across molecules^5^. We showed that the residual SNV detection rate of ppmSeq when applied to cell-free DNA was 3.5×10^−7^±7.5×10^−8^, comparable to duplex whole-genome sequencing^6^ and CODEC^11^.

Using *in vitro* and *in silico* mixing experiments, ppmSeq powered tumor-informed approach readily detected cancer across a range of mutational burden, as low as ∼1/Mb, which is typical for adult, solid tissue malignancies. Detection at plasma ctDNA fraction of 10^−6^ was achieved in cancer with high tumor mutation burden (>10/Mb) patients at 300x sequencing depth. These detection limits compare favorably to those obtained using Signatera, a widely used commercial panel for ctDNA detection with a limit of detection of 0.01%. Thus, for clinically relevant applications where input is extremely low, such as minimal residual disease detection after surgery and/or chemo-, radio- or immunotherapies, ppmSeq promises to be a transformative technology for sensitively monitoring disease recurrence in patients. Importantly, in plasma whole genome sequencing, breadth supplants depth of sequencing, thus libraries are far from fully saturated. As sequencing costs continue to decrease, deeper WGS of the plasma will become feasible, further enhancing essay sensitivity as shown in our forward-looking 300X mixing studies.

Currently, most methods for sensitive minimal residual disease detection require prior tumor sequencing and identification of targeted mutations in matched tumors^60–62^, limiting the ability to perform cancer monitoring in many clinical contexts. Excitingly, the low level of error that ppmSeq enables provides the opportunity to accurately identify disease even in the absence of matched tumor. As biopsy material is frequently scant and insufficient for sequencing^37,62^, there is an acute clinical need for such plasma-only tests. Towards this goal, we demonstrate that ppmSeq accurately detects ctDNA in the absence of matched sequencing information from the primary tumor for patients with urothelial and lung cancer, leveraging the trinucleotide contexts of genome-wide SNVs to detect ctDNA signal, which we showed tracks with imaging-based metrics.

We note that there are limitations to ppmSeq. Notably, as ppmSeq is a PCR-free library preparation method that relies on hydrogen bonds to carry both DNA strands into a single emulsion, it is not compatible with target enrichment methods relying on hybridization (and thus dsDNA denaturation), limiting it to a whole-genome assay. In addition, ppmSeq error rates are slightly higher than duplex sequencing methods, likely because traditional duplex sequencing relies on the consensus between two or more independently sequenced molecules to call variants, whereas dsDNA strands are sequenced from a single cluster in ppmSeq. The tradeoff between error rate and yield is therefore an important consideration and is likely to be application dependent.

Together, the high rate of dsDNA recovery and low error rates of ppmSeq provide a powerful tool for detection of ultra-low frequency variants. This has potentially major implications for an important future frontier of human genetics – somatic genetics. Recent groundbreaking work to genetically profile normal tissues has revealed extensive heterogeneity, where cells harbor distinct somatic variants resulting in vast clonal mosaicism even in heathy tissues^2,63–70^. To meet the challenges of profiling ultra rare somatic variants for complete picture of human mosaic tissue, we will need cost-efficient and accurate NGS solutions, such as ppmSeq. With such technology, the relationship between somatic mosaicism and human aging or non-malignant disease can be explored at unprecedented scale and resolution. Furthermore, in the liquid biopsy space, the ability of ppmSeq to deliver sensitive ctDNA detection through plasma WGS holds enormous promise for clinical application, including minimal residual disease monitoring after therapy and the potential for the development of early detection assays. We envision that ppmSeq will empower researchers and clinicians to ask new questions and design new assays to both advance our understanding of human genetics as well as push the boundaries for what is possible for disease detection and monitoring, to ultimately improve patient care.

## Acknowledgements

We thank the participants and their families for contributing plasma and tissue for this study. We also thank Henry R. He at Weill Cornell, Jane Park and all members of the laboratory of D.A.L., as well as the New York Genome Center computational biology team. A.J.W. received support from the Conquer Cancer Foundation Young Investigator Award, the Melanoma Research Alliance Young Investigator Award, and the NCI K08 Mentored Career Scientist Award. G.D.E. is supported by the National Institutes of Health Common Fund Somatic Mosaicism Across Human Tissues (UG3NS132024). MSKCC investigators are supported by Cancer Center Support Grant P30 CA08748 from the National Institutes of Health/National Cancer Institute. NYGC investigators are supported by the National Institutes of Health Common Fund Somatic Mosaicism Across Human Tissues (UM1DA058236). D.A.L. is supported by the Mark Foundation Emerging Leader Award, the Vallee Scholar Award, the Burroughs Wellcome Fund Career Award for Medical Scientists, a National Cancer Institute R01 grant (R01-CA266619-01), National Institutes of Health Common Fund Somatic Mosaicism Across Human Tissues (UG3NS132139) and the Melanoma Research Alliance Established Investigator Award. This work was made possible by the MacMillan Family Foundation and the MacMillan Center for the Study of the Non-Coding Cancer Genome at the New York Genome Center.

## Author Contributions

D.A.L., A.P.C. and I.R. conceived and designed the project.

A.J.W., M.A.A., M.S., J.M., A.K., D.W., J.O., O.E., J.M.M., M.S.M., J.E.S., G.D.E., B.M.F., A.Sa., M.K.C., N.R., A.R., S.Ge., and N.K.A. performed participant selection, curated participant data and prepared samples for sequencing.

A.G.T, R.F. and B.M.C. performed library preparation and sequencing.

A.P.C., I.R., A.So., E.M., G.K., O.H., D.S.T., S.Gi., A.J., O.B., S.A., S.R., T.P., D.J.Y., and D.A.L. performed data analysis.

A.P.C., I.R., E.M., G.K., O.H., C.P. and D.A.L. wrote the manuscript with comments and contributions from all authors.

## Code and Data Availability

Cell-free DNA alignment data and tumor-normal variant calls from the MIDGAM – Israel National Biobank for Research is available at https://www.ultimagenomics. com/resources/?category=datasets. Bioinformatic workflows for analyzing ppmSeq datasets are available at https://github.com/Ultimagen/healthomics-workflows/tree/v1.13.2. All other raw genomic sequencing data and clinical variables are being deposited to the European Genome-Phenome Archive. Code and custom scripts specific to the work presented here will be made available as a GitHub repository.

## Methods

### HUMAN SAMPLES

Blood and tissue samples were obtained from individuals in accordance with the Declaration of Helsinki. Samples were obtained from the Cryos International Sperm Bank (sperm sample), the Israel National Biobank for Research (tumor-informed analysis cohort), and the New York-Presbyterian/Weill Cornell Medical Center. Semen samples were obtained at Cryos International Sperm Bank from individuals enrolled in an IRB-approved protocol from New York University Grossman School of Medicine (IRB# i19-00794). Tumor, normal and plasma samples were obtained from MIDGAM -Israel National Biobank for Research (MID-116-2019), Weill Cornell Medicine (0201005295 (Tumor Biobanking), 1008011210 (GU Tumor Biobanking), 1011011386 (Urothelial Cancer Sequencing), 1007011157 (Genomic and Transcriptomic Profiling), 1305013903 (Precision Medicine), 1708018519 (Cardiac Surgery Biobank) and 1610017682 (ctDNA for Early Detection and Management of Non-Small Cell Lung Cancer).

### PPMSEQ DESIGN AND PROTOCOL

The ppmSeq adapters were designed for compatibility with the UG100 sequencer. The standard UG adapter sequences were extended by ppmSeq-specific sequences to include the AAAAAA bubble in both the start and the end of the single-end reads (in bold below). To enable adapter ligation without wasting excess sequencing cycles, each bubble is flanked by an 11bp dsDNA segments with a T overhang. The bubble sections are flanked by matched bases designed to ensure there is no cycle skip (see **Cycle-skip motifs in Ultima sequencing data** below) between the readouts of the two strands, as illustrated in the table below, where the measured homopolymer per flow is shown:

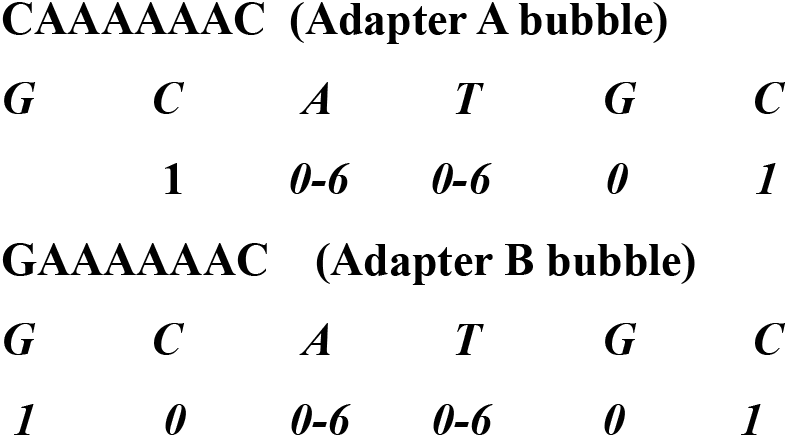

CAAAAAAC (Adapter A bubble)
|  |  |  |  |  |  |
| --- | --- | --- | --- | --- | --- |
| **G** | **C** | **A** | **T** | **G** | **C** |
|  | **1** | **0-6** | **0-6** | **0** | **1** |
| GAAAAAAC (Adapter B bubble) |  |  |  |  |  |
| **G** | **C** | **A** | **T** | **G** | **C** |
| **1** | **0** | **0-6** | **0-6** | **0** | **1** |

The ppmSeq adapters sequences are:

Adapter A:

~~~
5 ‘-ATCTCATCCCTGCGTGTCTCCGACTGCAC-
Barcode-GATC**AAAAAA**CATGAGCAGCAT-3’
~~~

And

~~~
5’-TGCTGCTCATG **AAAAAA** GATC-Barcode’-
GTGCAGTCGGAGACACGCAGGGATGAGATGGT-3’
~~~

Adapter B:

~~~
5’-
CATATGTGCGCG**AAAAAA**CATCACCGACTGCCCATAGAGA GCTGAGACTGCCAAGGCACACAGG-3’
~~~

And

~~~
5’-
AGCTCTCTATGGGCAGTCGGTGATG**AAAAAA**CGCGCACAT ATGT-3’
~~~

1.8–20 ng of cfDNA was used to generate PCR-free libraries using the KAPA HyperPlus PCR-Free Kit (KK8513; Roche). 1.5µL of ppmSeq adapters at concentration of 15µM of each ppmSeq adapters were used for ligation with the same adapter sequences above. Final library was cleaned using Agencourt AMPure XP beads for 1.2x followed by double-sided SPRI with ratios of 0.55x/1.2x. Sequencing was performed on a UG 100 single-end sequencer. For cfDNA samples, either the standard cfDNA library protocol (170 pM input) or the ppmSeq library protocol (105 pM input) was used.

To ensure that contaminating gDNA does not affect the cfDNA concentration measurement, we used a 4150 TapeStation System with a cfDNA tape to measure the cfDNA concentration by quantifying the sample concentration in the 100-700 bp size-range. Finally, we used a quantitative PCR (qPCR) assay to quantify concentrations of PCR-free DNA libraries for sequencing on the UG 100™ platform using NEBNext^®^ Library Quant Kit for Ultima Genomics (E3410).

### DNA EXTRACTION AND SEQUENCING

#### Sperm DNA sample

Sperm purification and DNA extraction was performed according to Lui et al^27^. Genomic DNA underwent blunt-end fragmentation using the HpyCH4V restriction enzyme, followed by a 2.2× AMPure bead cleanup, nick ligation with E. Coli DNA Ligase (NEB), a 2.2× bead cleanup, A-tailing with a dATP/ddBTP mix, and a 2.2× AMPure bead cleanup according to Liu et al^27^. The resulting DNA was used to generate ppmSeq PCR-free libraries using the KAPA HyperPlus PCR-Free Kit (KK8513; Roche), following the manufacturer’s protocol and per the above ppmSeq adaptor ligation protocol. For ligation, 2.5 µL of ppmSeq adapters at a concentration of 15 µM each were used. The final libraries were cleaned using Agencourt AMPure XP beads with a 1.1× cleanup, followed by a double-sided SPRI cleanup (0.65× / 1.1×). 105 pM of the final library was used for single-end sequencing on a UG 100 sequencer.

#### Tumor-informed analysis samples from MIDGAM – Israel National Biobank for Research samples

Fresh Frozen was extracted using the AllPrep DNA/RNA Mini Kit (20-80204; Qiagen). PBMC DNA was isolated with the DNeasy Blood & Tissue Kit (20-69504; Qiagen). PCR-free sequencing libraries for both Fresh Frozen and PBMC samples were generated from 200 ng of DNA using the KAPA HyperPlus PCR-Free Kit (KK8513; Roche) after Covaris shearing.

Plasma was collected in Greiner’s Vacuette tubes Cat# 455036 (Vacuette Tube 9ml with K3EDTA Greiner Bio-One). cfDNA was extracted from 4 mL of plasma using the cfPure® V2 Cell-Free DNA Extraction Kit (PN: K5011610-V2, BioChain). For standard cfDNA sequencing, libraries were prepared from 4.2–10 ng of input using the xGen Prism DNA Library Prep Kit (IDT). For ppmSeq, 3.7–20 ng of cfDNA was used to generate PCR-free libraries using the KAPA HyperPlus PCR-Free Kit (KK8513; Roche). 1.5µL of ppmSeq adapters at concentration of 15µM of each ppmSeq adapters were used for ligation with the same adapter sequences above. Final libraries were cleaned using Agencourt AMPure XP beads for 1.2x followed by double-sided SPRI at ratios of 0.55x/1.2x. Sequencing was performed on a UG 100 single-end sequencer. For cfDNA samples, either the standard cfDNA library protocol (170 pM input) or the ppmSeq library protocol (105 pM input) was used, achieving average coverage depths of 138x and 70x, respectively.

#### Urothelial cancer, cancer-free control and lung cancer samples

Urothelial cancer somatic mutations obtained from tumor-normal sequencing were obtained from Nguyen et al^39^. UMI-duplex sequencing data from cancer-free controls and urothelial cancer patients were previously published^6^, and remaining cell-free DNA from these samples was used to generate ppmSeq libraries for this study. Cell-free DNA from all three cohorts was extracted from plasma using the Magbind ccfDNA extraction kit (K2EDTA or Streck, Omega Biotek, M3298). Manufacturer recommendations for extraction were followed, but elution volume was set to 35uL, and elution time was increased to 20 minutes on a thermomixer set to 1,600rpm at room temperature. Extracted DNA was quantified using a Qubit 3.0 fluorometer and sequenced on a UG 100 sequencer using 105pM of ppmSeq library.

### BIOINFORMATIC ANALYSIS OF PPMSEQ DATA

#### Cycle-skip motifs in Ultima sequencing data

Ultima sequencing’s flow-based sequencing by synthesis technology cycles through flows of non-terminated nucleotides in a repetitive cycles (T,G,C,A) and reads the length of each incorporated homopolymer to generate sequencing reads, which are measured in flow-space and converted to base space^12^ while error probabilities are reported in flow-space^71^. Sequences differing by one substitution in base-space differ either by 2 “flows” (homopolymers measured on different bases), or more – some substitutions termed cycle skip motifs cause a shift in the number of sequencing cycles compared to the reference^6^ (**Supplementary Table 12**).

#### Identification of dsDNA-derived sequencing reads in ppmSeq

During the ppmSeq workflow, the bubble-adapter templates in the start and end of the insert are sequenced. In reads comprised of both original strands (which we refer to as “mixed”), equal proportions of A and T sequencing signals are expected in dsDNA-derived sequencing reads, whereas A or T sequencing signals are expected if one of the strands dropped out. Therefore, we empirically defined mixed reads as sequencing reads with A/(A+T) sequencing ratios between 0.27 and 0.73 in both adapter positions, (allowing calls of 2A5T or 5A2T but not 2A6T or 6A2T) and a total called homopolymer length between 4 and 8 (expected value is 6, see **Supplementary Figure 1** for A/ (A+T) ratios). Reads harboring calls of polyA or polyT only in the same total homopolymer range were annotated as plus- or minus-strand-only reads, respectively (**Supplementary Figure 1**). These categorical annotations were used as part of the SNV quality recalibration method used for the tumor informed analysis, while only mixed reads were used in the tumor-naive analysis which is more sensitive to background noise.

#### SNV denoising and quality recalibration

Our read-centric de-noising framework was developed to overcome the limitations of traditional locus-centric variant calling, particularly in scenarios where rare mutations may be supported by only a single read. To achieve this, we generated a comprehensive dataset capturing every candidate SNV along with a rich suite of annotations that describe sequencing quality, local sequence motifs, and fragment-specific features, detailed below. Among all SNVs in high-coverage (≥20×) regions, those supported by multiple reads (variant allele frequency greater than 0.7, indicating a high likelihood of representing genuine homozygous germline variants) were labelled as True, while False SNVs were defined as singleton events (SNVs with a single read support, indicating they are more likely to be artifacts). Using these curated sets, we trained an XGBoost^72^ classifier to robustly distinguish between true and artifactual SNVs. Once trained, the classifier assigns a calibrated quality score to each SNV, providing a precise estimate of the residual error rate. For the tumor informed analysis, a model was trained for each sample separately. For the tumor-naive analysis, only mixed reads were retained, and samples were batched together to increase the number of artifactual SNVs available for model training. Given that the sperm DNA sample was sequenced on its own wafer and had achieved high coverage (100x), a model was trained separately.

We now provide detailed information on the denoising procedure outlined above.

##### Training set preparation

Artefactual SNVs can derive from sequencing errors as ppmSeq does not use single strand UMI based denoising or from library error, especially due to overhangs that are not removed in cfDNA libraries as fragment blunting methods are not compatible with cfDNA and would have a matching variant on both strands after end repair. To filter out artefactual SNVs, we employed a supervised machine learning model trained to classify actual SNVs (labelled True or TP) form noise (False or FP). The dataset preparation pipeline consists of the following steps.

##### SNV extraction

Only reads with a mapping quality of 60 (maximal value in the aligner) are considered. From these reads, all SNVs with respect to the reference genome, where the adjacent bases () from the respective SNV match the reference genome, are extracted to a separate VCF file. The adjacent base filter is intended to avoid calling compound homopolymer errors as SNVs.

##### Data labeling

A labeled dataset is created as follows. All reads that support a SNV with respect to the reference are examined by creating a pileup. Only pileups with coverage ≥20× are considered. When > 70% of the reads in the pileup support the variation, the SNV is labelled ‘True’. When only a single read supports the SNV while the rest support the reference, the SNV is labelled as ‘False’. All other reads are excluded from training.

##### Genomic region filtering

Training data was filtered for SNVs included in the Ultima Genomics High Confidence Region in chromosomes 1-22. The UG high-confidence region (HCR) covers >99% of the GIAB v4.2.1 HCR and excludes genomic areas where UG performance is consistently of lower confidence, such as homopolymer regions of length >12 bp and limited regions of low complexity (see https://github.com/Ultimagen/healthomics-workflows/blob/v1.13.2/docs/ug_hcr.md). Additionally, SNVs represented in dbsnp^73^ or in gnomAD^74^ with AF>0.001 were excluded to avoid risk of contamination introducing labeling noise. When tumor sequencing was available for a patient ctDNA sample, SNVs intersecting the tumor mutations were also excluded from the training set preparation.

##### Feature extraction

The features used to characterize the SNVs include local and read-wise quality scores, sequence motif features, and information about reads. Boolean and categorical features are explicitly indicated below; all other features are numerical. For more details refer to **Supplementary Table 2**.

##### Sequencing quality features

The read Ultima qualities are translated to several quality-related features per called SNV: is_cycle_skip is a Boolean feature indicating whether the SNV entails a cycle-skip with respect to the reference: X_ SMQ_LEFT and X_SMQ_RIGHT measure the median base quality of the 20 bases to the left/right of the SNV. Finally, rq is the read quality score in the first 100 flows where Q is the per flow quality Phred score (equation 1).

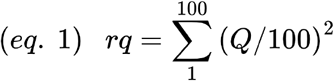

##### Edit distance

X_EDIST, X_FC1, X_FC2 are three measures of the overall edit distance between read and reference – X_EDIST is the Levenshtein distance from the reference and includes SNVs and indels, X_FC1 is the total number of SNVs, and X_FC2 is the total number of SNVs where the adjacent bases () from the respective SNV match the reference genome.

##### Sequence motif features

ref and alt are the identities of the reference and alternate nucleotides, and prev3, prev2, prev1, next1, next2, next3 are the immediate 3-base context flanking the variant (in order, in the order of the reference genome (e.g. next1 is the next base in the reference genome in the forward orientation). All of these are categorical features (with values A, C, G, or T). hmer_ context_ref/hmer_context_alt are the lengths of the homopolymers that contain the SNV base in the ref/alt allele.

##### Information about the read

strand_ratio_cagtegory_ start / strand_ratio_category_end are categorical features containing information about the ppmSeq tags at the start/end of the read. For the convenience of discussion, when both equal “MIXED” we refer to the read as mixed, otherwise we consider it to be non-mixed (but note that this Boolean mixed/non-mixed information is not a feature used for model training, rather the model is provided the richer information of the ppmSeq tags each end). X_ LENGTH, X_INDEX are the length of the insert and the relative position of the SNV in it. is_forward is a Boolean indicating the read direction. max_softclip_length is the maximal softclip length at either end of the read.

Roughly speaking, a cycle-skip SNV (which causes a shift in all cycles following the SNV) is with high probability free of sequencing errors, whereas SNVs with mixed ppmSeq tags are with high probability free of errors due to library preparation artifacts. It might thus be expected that these should be the two most consequential features in determining the quality of an SNV. In fact, a SHAP-score^75^ based examination of feature importance (**Supplementary Figure 7C**) reveals that the contributions to the model prediction of all the features are of the same order of magnitude (for example, the sum of SHAP scores of the 164 least impactful features is on average larger than the SHAP score of the most impactful feature. Note that the ppmSeq tag features are absent from this plot as the pre-filter for the presented data retains only mixed SNVs, as described next).

The distributions of feature values by label are presented in **Supplementary Figures 5** and **6** and **Supplementary Table 5**.

##### Pre-filtering

Only mixed reads with EDIST<10, X_ LENGTH<220 and Base Calling Sequencing Quality > 7.5 are retained, the others are excluded from training.

##### Downsampling

To ensure that sequence features of homozygous germline SNVs do not bias model predictions (for example prioritizing C>T transitions that would not be meaningful for all studies), the True dataset was down-sampled so that all trinucleotide contexts (including the alt base, e.g. ACG>ATG), have equal abundance. Forward and reverse complement motifs were counted separately to form distributions with 192 motifs, because while biological signals are symmetric to reverse complement, sequencing artifacts are generally not. The resulting uniform histograms of ‘True’ SNVs by trinucleotide context is presented in **Supplementary Figure 3A**. Next, both ‘True’ and ‘False’ SNVs were down-sampled to a dataset of no more than 1.5M SNVs of each label (the resulting training set might be unbalanced when less than 1.5M SNVs of either label is available).

##### Model

We trained XGBoost models with log-loss on the labelled dataset described above, using the hyperparameters are described in **Supplementary Table 3** (model performance did not sensitively depend on hyperparameter values). We employ a 5-fold cross-validation (CV) design where the data is split into 5 folds by chromosomes, grouped into 5 comparable size groups using a greedy load balancing algorithm. CV by position ensures that there is no data leakage between the training set and the predictions of the model at inference time.

##### Model evaluation

As explained above, we employ a 5-fold cross-validation design. To conduct a meaningful comparison of model performance on datasets with different base rates (fractions of True/False SNVs), we measure model performance by the ROC AUC score (which does not explicitly depend on the base-rate), unless otherwise indicated. ROC AUC scores for models trained on samples from the Control, Lung, and Bladder batches are presented in **Supplementary Figure 5A**. As can be seen, model performance improves when the training sets are larger and more balanced. To obtain larger training sets we pool data from different samples, to train both batch-level models (a single model for each batch), and a multi-batch model (a single model for all batches. **Supplementary Figure 5B** shows that the batch-models achieve highest ROC AUC scores. These are the models we employ in the tumor-naive analyses throughout the paper.

To obtain further insight into how these features inform the model, we study the SHAP (SHapley Additive exPlanations) values of 3 batch-level models, see **Supplementary Figure 5C**. The SHAP score indicate that X_SCORE and SNV trinucleotide context consistently have the highest SHAP scores, but many of the features contribute to the models’ outputs.

##### Quality recalibration, SNVQ metric, and XGBoost-based filtering

Given a filter that selects a subset of all SNVs, we measure its denoising quality by Filter Quality (FQ), defined as:

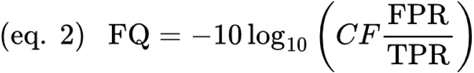

Where FPR (false positive rate) and TPR (true positive rate) are measured as the fraction of all SNVs in the test set labelled respectively FP and TP that pass the filter. CF is a calibration factor that takes into account how data was selected for the test set:

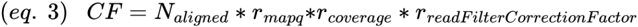

Where *r*_*mapq*_ is the fraction of reads with mapq=60 (maximal value allowed in the aligner), *r*_*coverage*_ is the fraction of reads in regions where the coverage is at least 20 and at most the 95^th^ coverage percentile, and *r*_*read_filter_correction_factor*_ is the fraction of the reads in the TP SNV loci that are filtered due to adjacent differences from the reference genome, and

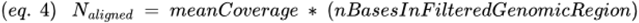

is the effective number of bases used in the analysis.

Note that the FQ score is related to the precision of the filter through the *base rate*, the fraction of all True SNVs:

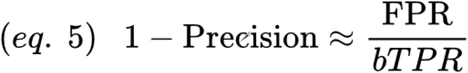

Where the approximation holds when

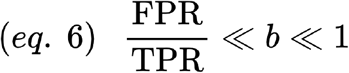

Expressed in Phred scale:

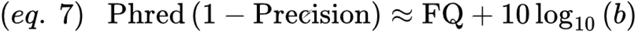

The XGBoost model estimates for every SNV the probability that it is True given the feature values. This estimate can be used to filter SNVs by retaining only those with estimated probabilities above a threshold. This filtering strategy defines a calibration function that maps the threshold probability value to the FQ of the resulting filter. We use this calibration function to assign a quality score SNVQ to every SNV: the XGBoost model is applied to the SNV features to produce an estimated probability, and this probability is pushed forward through the calibration function to produce the SNVQ. In other words, SNVQ is the FQ of the filter whose threshold is the model’s prediction for the SNV in question.

#### Genome In a Bottle-mixture experiment

Genomic DNA of HG001 and HG005 were in-vitro diluted into HG002 gDNA in 0, 1, 10 and 100 parts-per-million fractions. ppmSeq libraries were sequenced to a 30x average coverage. To estimate GIAB fraction, we generated two lists of HG001- and HG005-unique homozygous SNVs, using the Broad GIAB v4.2.1 ground-truth variants and high confidence regions (HCRs)^76^. We limited the variants to SNVs found inside the intersection of HCRs of HG001, HG002 and HG005, and are not adjacent to another GIAB’s variant within two base-pairs (bedtools slop -b 2). To avoid the detection of rare variants which are not in the ground truth, we generated a pileup of HG002 variants inside the HCRs, based on a HG002 sample we sequenced independently to this experiment. We then filtered out rare variants represented by two or more reads. Final signatures contained 77944 SNVs (HG001) and 130698 SNVs (HG005). The number of variants per filtering stage are listed in **Supplementary Table 13**. ppmSeq reads underwent SNV denoising and quality recalibration (**Methods**), resulting in SNV quality score (quality) per read. Reads with quality>62 where both tags were Mixed were further intersected with GIAB signatures. To evaluate accuracy and match a number of variants that is more typically seen in cancer genomes, we subsampled GIAB-unique signatures to a size of 30,000 SNVs for 30 repeats. Bootstrapping results are listed in **Supplementary Table 5**.

#### Tumor-informed ctDNA detection through in silico mixing studies

To evaluate the limit of detection (LoD) of our ctDNA detection approach, we performed in-silico mixing studies that simulate MRD detection across a range of tumor fractions. In these analyses, MRD-positive samples with known tumor fractions were computationally mixed with reads from control samples (cross-patient controls obtained by intersecting tumor variants from one patient with cfDNA from another or healthy individual controls). Supporting mixed reads from both matched and control individuals were first extracted and filtered based on a quality threshold of SNVQ>62 —derived from the quality recalibration algorithm described above—and then intersected with tumor-derived somatic mutation calls. For a series of predefined tumor fraction values, we simulated the expected number of supporting reads using a Poisson distribution model, thereby accounting for differences in sequencing coverage between samples. In each realization simulating a simulated coverage *cov*_*sim*_ from a sample with a coverage *cov*_*original*_ where *n*_*original*_ background reads (intersection between a patient mutation signature and a healthy or cross-patient plasma sample) were measured, we simulated the number of background reads in the target coverage using

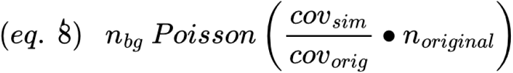

and the number of signal reads for a simulated tumor fraction *TF*_*sim*_ from an original tumor fraction *TF*_*original*_ using

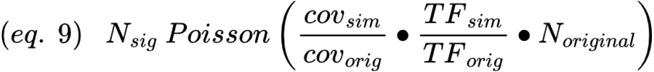

where *N*_*original*_ is the number of signal reads (intersection between a patient mutation signature and its plasma). For each simulated condition, we drew 100,000 samples and computed a z-score as the difference between the matched and background read distributions normalized by the background’s standard deviation. The aggregated results across control patients yielded a calibrated measure of ctDNA detectability, informing the threshold at which true tumor-derived signals can be confidently distinguished from sequencing artifacts. We performed this procedure 10 times per condition (n = 50 replicates per condition).

#### De novo mutational identification (sperm DNA sample) and plasma-only ctDNA detection using mutational signatures

To further improve the specificity of mutation detection, SNVs were denoised based on a cohort-level quality score cutoff that was derived for each trinucleotide (see ***Trinucleotide motif-specific thresholding***). The quality scores for True variants (i.e., germline variants) and False variants (single supporting reads) were compared for each 192 trinucleotides, and a cutoff was selected according to Younden’s index or 12, whichever was higher. Next, an allelic frequency cutoff of 0.2 was used to remove germline variants (except for data presented in **Figure 3D**, where normal sequencing data was used to remove germline variants). To measure mutational signatures, an Ultima-specific signature was derived by tabulating variants from a set of 20 cancer-free controls. Urothelial cancer patient frequencies were fit to a catalog of SBS2, SBS13, SBS31, SBS35, clonal hematopoiesis^41^ and the Ultima-specific signature using the MuSiCal signature fitting tool^77^. APOBEC3-specific signal was calculated by summing the contribution of SBS2 and SBS13, and SBS31 alone was used to infer platinum chemotherapy-specific signal, as we found SBS35 to be incorrectly fit and cause false positive signals in cancer-free controls. Lung cancer cfDNA samples were fit to a custom catalog of SBS4, clonal hematopoiesis and the Ultima-specific signature. Cancer-free controls were fit to the urothelial cancer catalog or the lung cancer catalog, depending on the analysis. Importantly, the cancer-free controls used to derive the Ultima-specific signature were not used during catalog fitting. Performance of plasma-only ctDNA detection using mutational signatures was assessed using ROC analysis comparing cases (ie, a cohort of a certain cancer type) to cancer-free controls fit to the same catalog).

#### Trinucleotide motif-specific thresholding

We have previously reported that Ultima sequencing’s flow-based technology has trinucleotide-specific error rates that are determined based on the specific order by which the bases are added during sequencing (traditionally T,G,C,A)^6^. Therefore, we reasoned that trinucleotide-specific thresholds might enable denoising that maximizes signal-to-noise ratios without introducing trinucleotide biases. To accomplish this, SNVQ thresholds were selected per trinucleotide motif. Optimal cutoffs were selected according to Youden’s index. Importantly, when tabulating trinucleotide frequencies using the commonly used format of collapsing trinucleotides according to central purines (96 trinucleotide format), we observed different quality distributions that corresponded to whether a central purine or central pyrimidine was reported in the original sequencing read. Therefore, cutoffs were determined and applied by considering the trinucleotide as it was sequenced, without collapsing the pyrimidines with purines.

### COMPARING DOUBLE-STRANDED DNA RECOVERY IN PPMSEQ, NANOSEQ AND CODEC DATASETS

#### ppmSeq datasets

ppmSeq dsDNA yields were defined as the number of mixed reads (as defined above) divided by the total number of sequenced reads.

#### NanoSeq datasets

NanoSeq yields prior to filtering were obtained from Abascal et al. (their Supplementary Table 2). Briefly, their data was subset to only contain NanoSeq libraries, and yields prior to denoising were estimated by dividing the number of read bundles containing two or more top strands and two or more bottom strands, and dividing by the total number of paired-end reads sequenced. After denoising, the residual SNV rate was taken from their Supplementary Table 4 (“burden”) column, and the dsDNA recovery after filtering was estimated by dividing the number of sites called (their Supplementary Table 2) divided by the number of bases sequenced (their Supplementary Table 2, “PE reads sequenced” multiplied by the bases sequenced per read pair (300)).

#### CODEC datasets

Data were obtained from Bae et al. (their Supplementary Table 1: CODEC WGS) and CODEC yields prior to filtering were estimated according to:

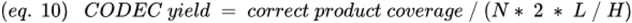

Where *N* represents the number of paired-end sequenced reads, *L* represents the read length, and *H* represents the size of the human genome used for alignment (hg19; 2,897,310,462 non-N bases). The denoised yield was estimated by dividing the final number of duplex bases (their Supplementary Table 1) by the number of bases sequenced (the number of reads (2 times the number of read pairs), multiplied by the read length).

#### Statistics and plots

All statistics were calculated in R (version 4.4.1) or python (version 3.6). Boxplots were generated using the ggplot2 R package or matplotlib python package. The lower and upper ends of the boxes represent the 25th and 75th percentiles of the data, respectively, and the horizontal line represents the median. The whiskers represent at most 1.5 times the IQR.

## SUPPLEMENTARY FIGURES

**Supplementary Figure 1.**
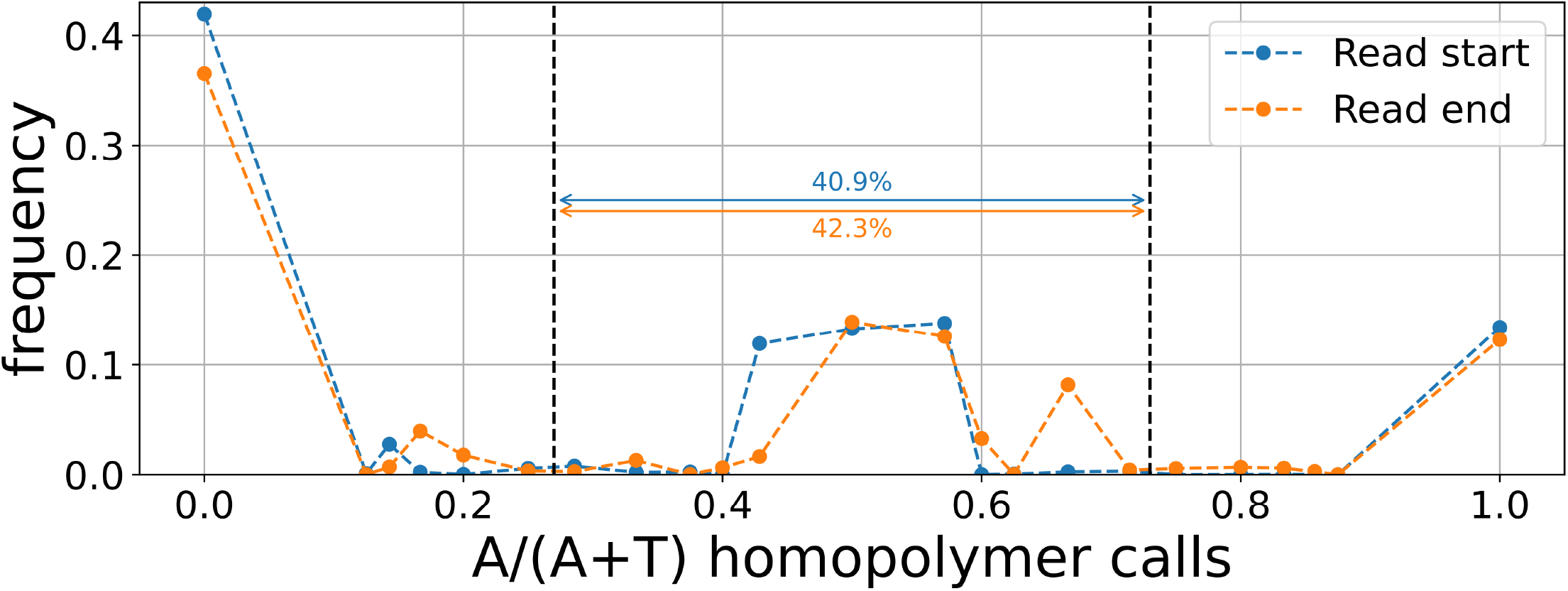
Histogram of homopolymer calls in sample UPN-001. This plot is a histogram of the ratio of A homopolymer to T homopolymer in either the bubble section in the start (blue) or end (orange) of the read, with the limits of [0.27,0.73] annotated (dashed black lines). Total percent of mixed reads are shown as text.

**Supplementary Figure 2.**
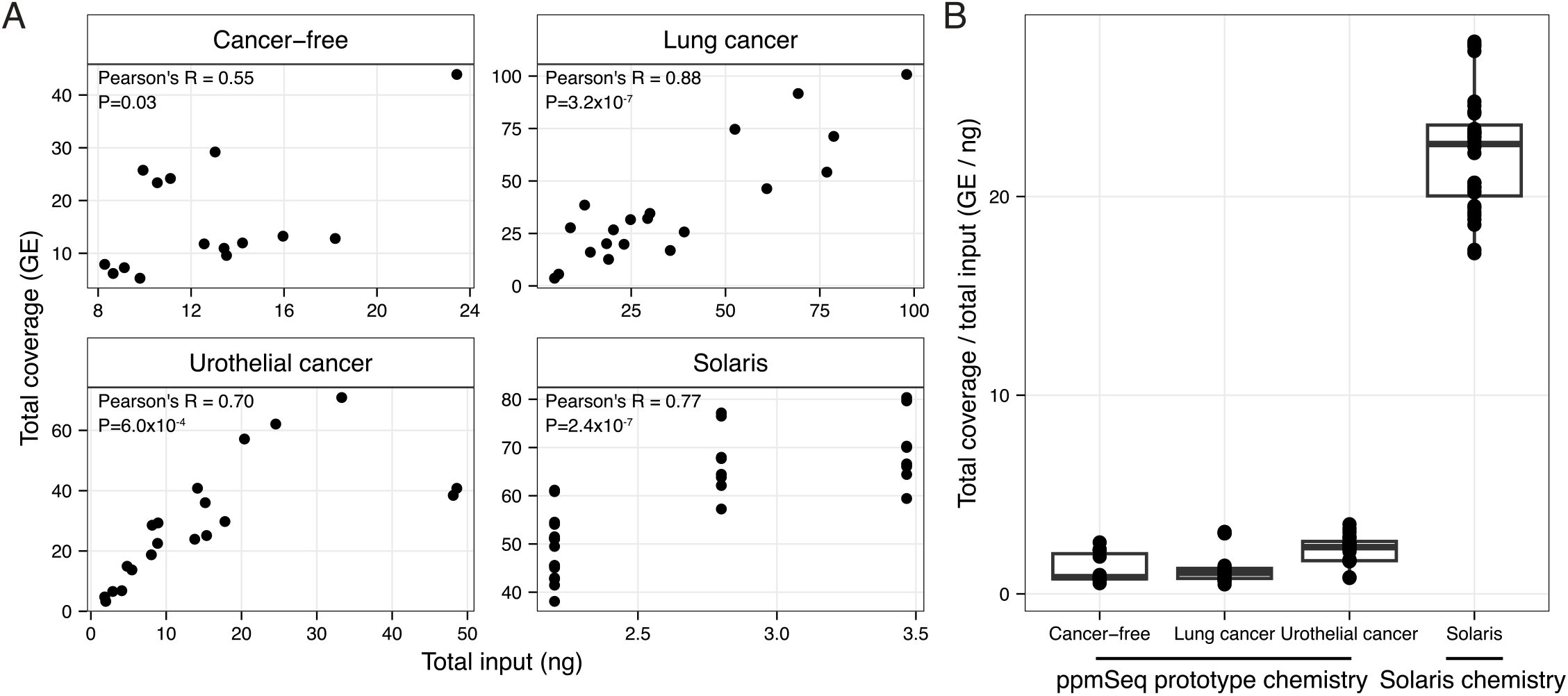
ppmSeq coverage yields in different batches. **A** Assessment of total genome coverage [genome equivalents (GE)] across different total input mounts (ng) of cell-free DNA (cfDNA) sequenced using ppmSeq in four different batches: cancer-free controls, patients with lung cancer, patients with urothelial cancer and samples prepared with the commercially available Solaris workflow. **B** Coverage-to-input ratios of the four sequencing batches. Ratios were obtained by dividing the total coverage obtained using the ppmSeq protocol by the library input, in nanograms.

**Supplementary Figure 3.**
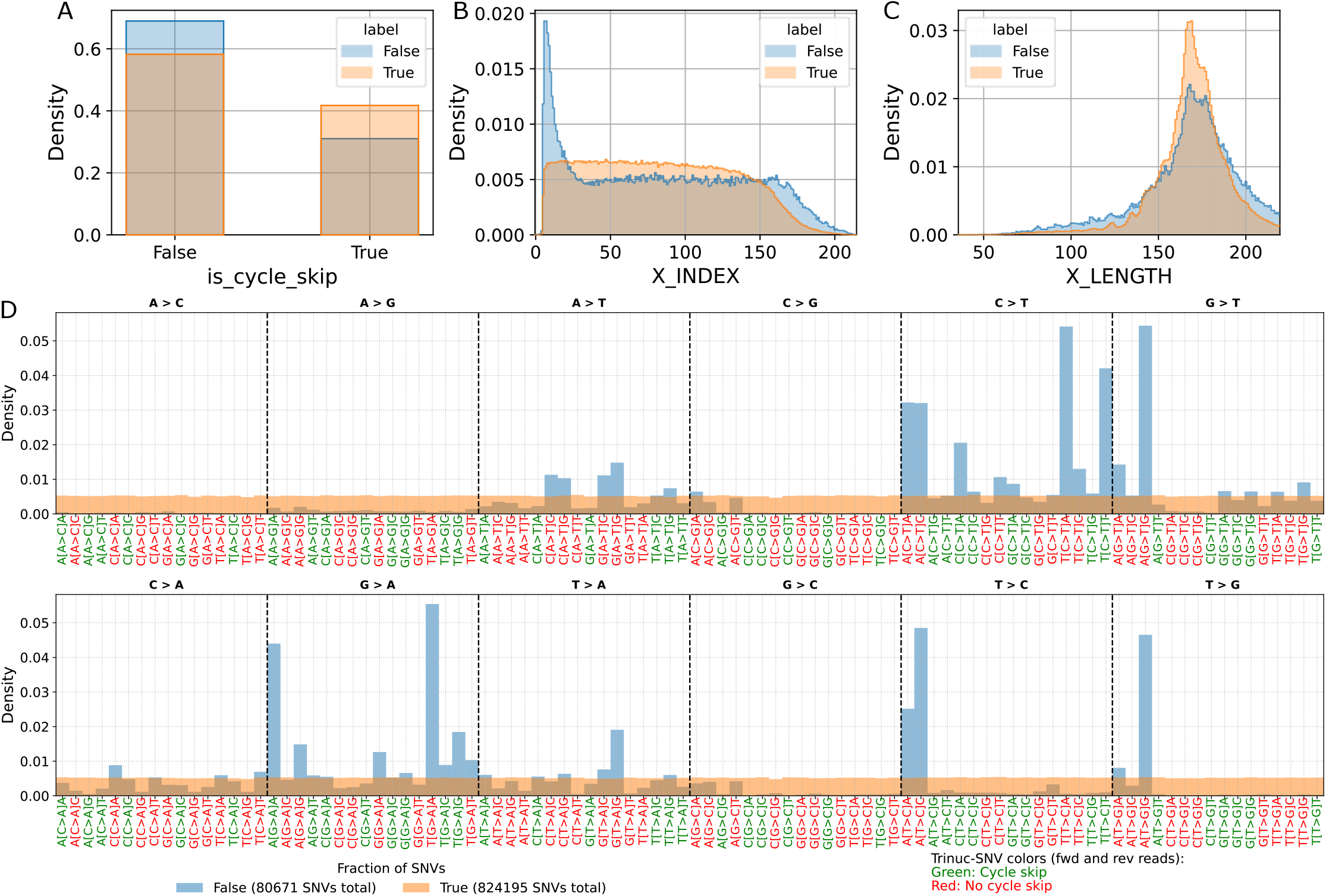
The distribution of feature values in the training set for a few select features, for False (single-supporting read) and True (Homozygous germline) SNVs from the cancer-free controls. **A** When a reported trinucleotide mutation would cause a shift in the number of sequencing cycles compared to the reference, the trinucleotide context is more robust to sequencing errors^6^ (**Methods**). is_cycle_skip is ‘True’ for contexts where such a shift occurs, and False otherwise. **B** X_INDEX is the position along the read, demonstrating an increase in artifacts near the read start. These errors represent errors on single-stranded overhangs that are copied onto the other complimentary during end-repair^6,8,28^ **C** X_LENGTH is the insert length, as longer cfDNA fragments have been shown to have a higher proportion of nicked DNA, which can propagate DNA damage-induced errors^8,76^. **D** probability as a function of the SNV trinucleotide context and alt values. The green/red colors of SNV context labels indicate cycle-skip/non-cycle-skip SNVs respectively. To ensure that sequence features of homozygous germline SNVs do not bias model predictions (for example prioritizing C>T transitions that would not be meaningful for all studies), the True dataset was down-sampled so that all trinucleotide contexts (including the alt base, e.g. ACG>ATG), have equal abundance. Forward and reverse complement motifs were counted separately to form distributions with 192 motifs, because while biological signals are symmetric to reverse complement, sequencing artifacts are generally not.

**Supplementary Figure 4.**
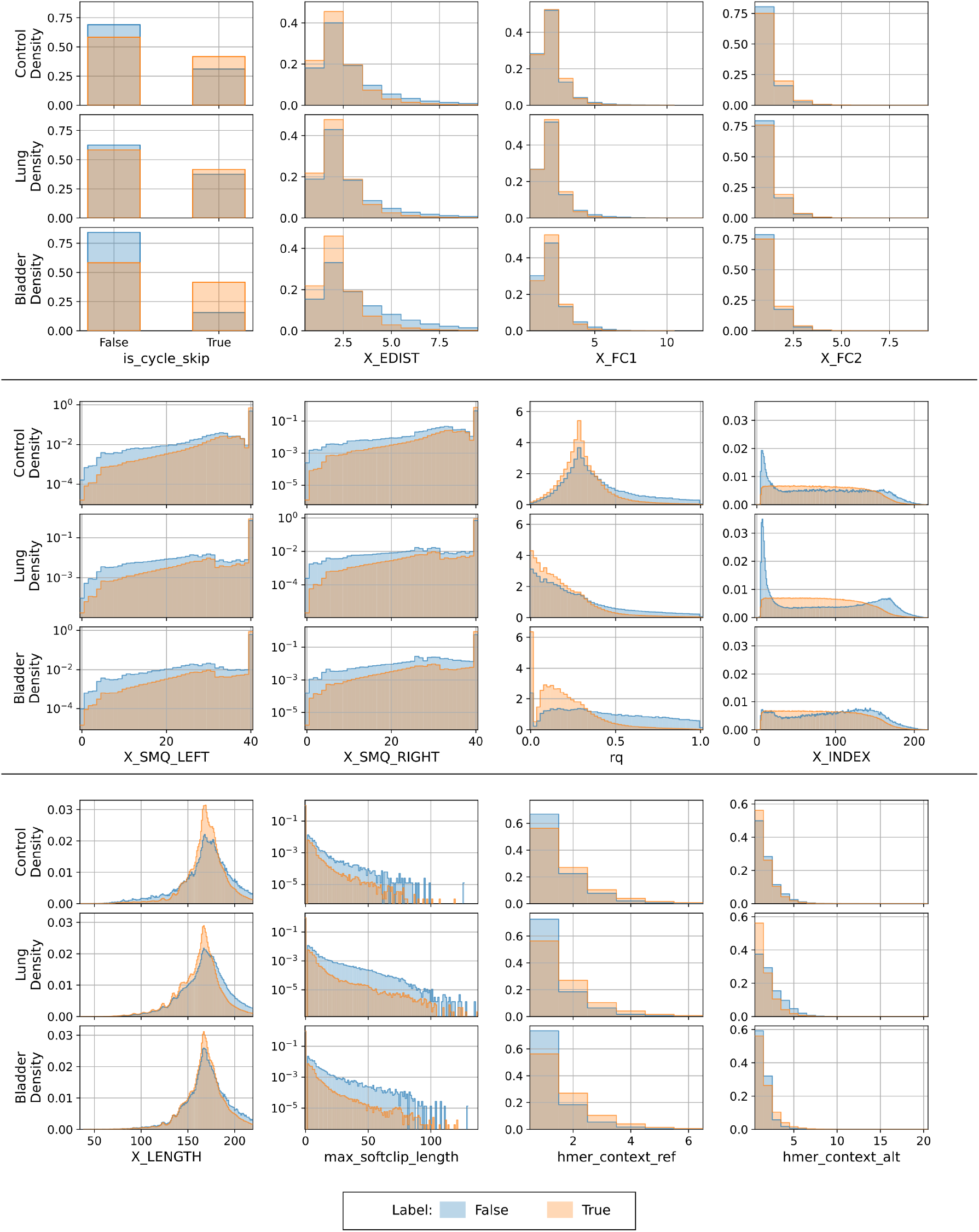
Histograms for 12 features for Control, Lung and Bladder batches, normalized separately over the ‘True’ and ‘False’ SNVs in the training sets. Quality-related features: is_cycle_skip is a Boolean feature indicating whether the SNV entails a cycle-skip with respect to the reference. X_EDIST, X_FC1, X_FC2 are three measures of the overall edit distance between read and reference – X_EDIST is the Levenshtein distance from the reference and includes SNVs and indels, X_FC1 is the total number of SNVs, and X_FC2 is the total number of SNVs where the adjacent bases (±5bp) from the respective SNV match the reference genome. X_SMQ_LEFT and X_SMQ_RIGHT measure the median base quality of the 20 bases to the left/right of the SNV. Finally, rq is the read quality score, defined as rq= in the first 100 flows where Q is the per flow quality Phred score. Information about the read: X_LENGTH, X_INDEX are the length of the insert and the relative position of the SNV in it. max_softclip_length is the maximal softclip length at either end of the read. Sequence motif features: hmer_context_ref/hmer_context_alt are the lengths of the homopolymers that contain the SNV base in the ref/alt allele. Further information about all features is detailed in **Supplementary Table 5**.

**Supplementary Figure 5.**
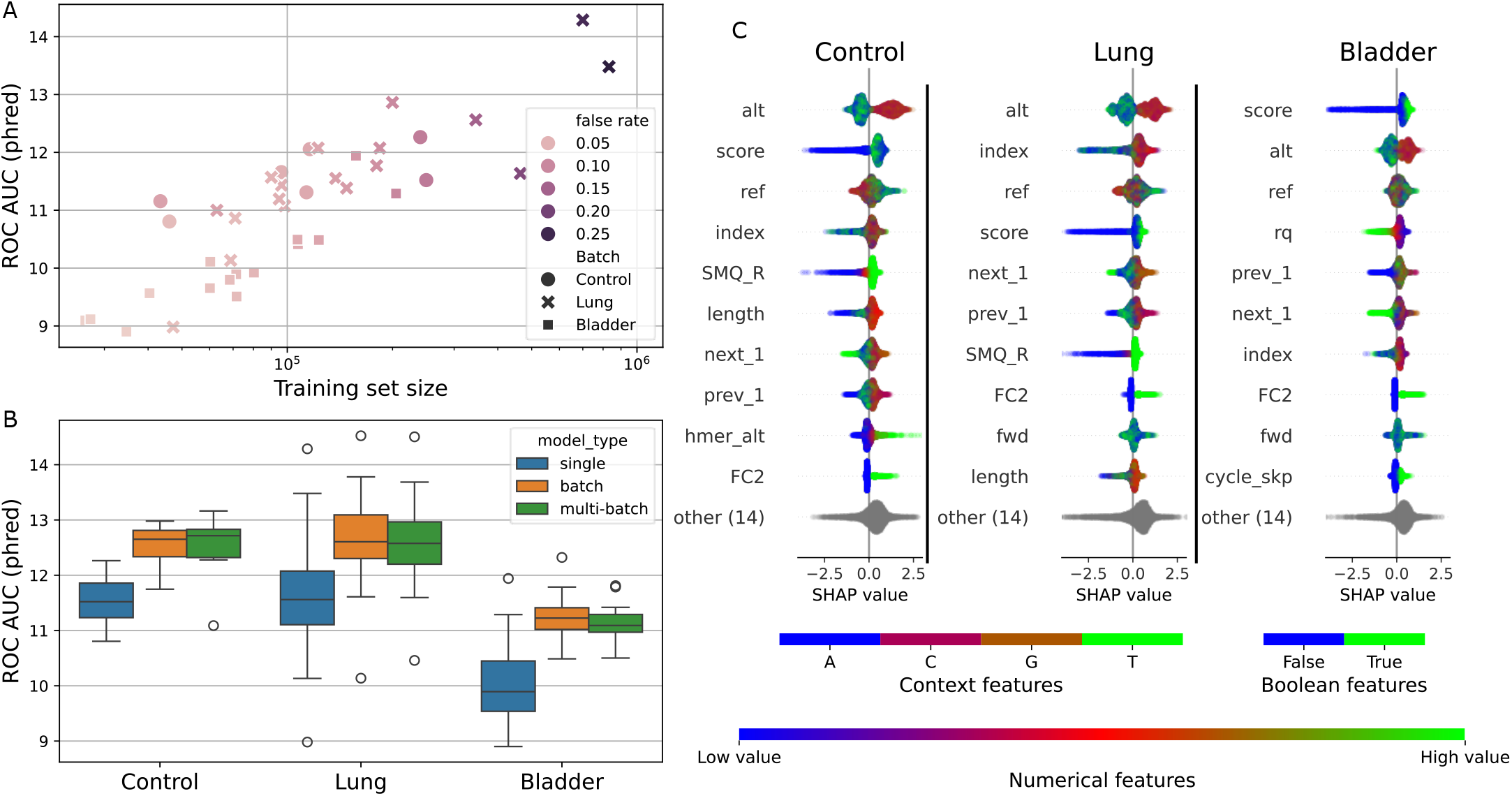
XGBoost model metrics by cohort. **A** Trained XGBoost model performance for 40 single-sample models, as a function of the training set size and the False rate (fraction of training SNVs with label ‘False’). Model performance here is measured by the ROC AUC for the out-of-fold model predictions compared with the SNV label. These training sets were steeply pre-filtered to retain only high-quality (X_SCORE>7.5) mixed SNVs; sequencing datasets that contain relatively few pre-filtered False SNVs produce smaller and more imbalanced training sets, and display poorer model performance. **B** The 40 samples belong to 3 cfDNA batches: Control (7 samples), Lung (18), and Bladder (15) (see **Supplementary Table 1** for sample-to-batch key). To overcome the scarcity of False SNVs in the training data, samples were pooled to create 3 batch level training sets comprised of data from all samples belonging to each of the 3 batches, and also to a multi-batch training set comprised of data from all 40 samples. The resulting batch and multi-batch level models were found to perform better than single-sample models. Subsequently, single-sample models were used for the tumor informed analysis, where both mixed and non-mixed reads were retained. In the tumor-naive analysis, only mixed reads were retained, and batch-level models were used. **C** SHAP (SHapley Additive exPlanations^73^) feature importance scores for the three batch-level models. Colors indicate feature values. Features are listed in descending order of importance, as measured by mean absolute SHAP score value. “other” is the sum of the other 14 features not presented in the figure. In addition to the features presented in Supplementary Figure 6, these include: ref/alt, the reference and alternate bases of the SNV; prev3, prev2, prev1, next1, next2, next3 are the immediate 3-base context flanking the variant (in order), in the order of the reference genome (e.g. next1 is the next base in the reference genome in the forward orientation); X_SCORE, a base-quality-like score for the SNV, defined as the log likelihood difference between the base-qualities of the ref and alt sequences in flow-space. For clarity of the figure, feature names were shortened: X_ SCORE to score, X_LENGTH to length, X_INDEX to index, X_SMQ_RIGHT/LEFT to SMQ_R/L, is_forward to fwd, is_cycle_skip to cycle_skp, hmer_context_ref/alt to hmer_ref/alt. Further information about the features is detailed **Supplementary Table 5**.

**Supplementary Figure 6.**
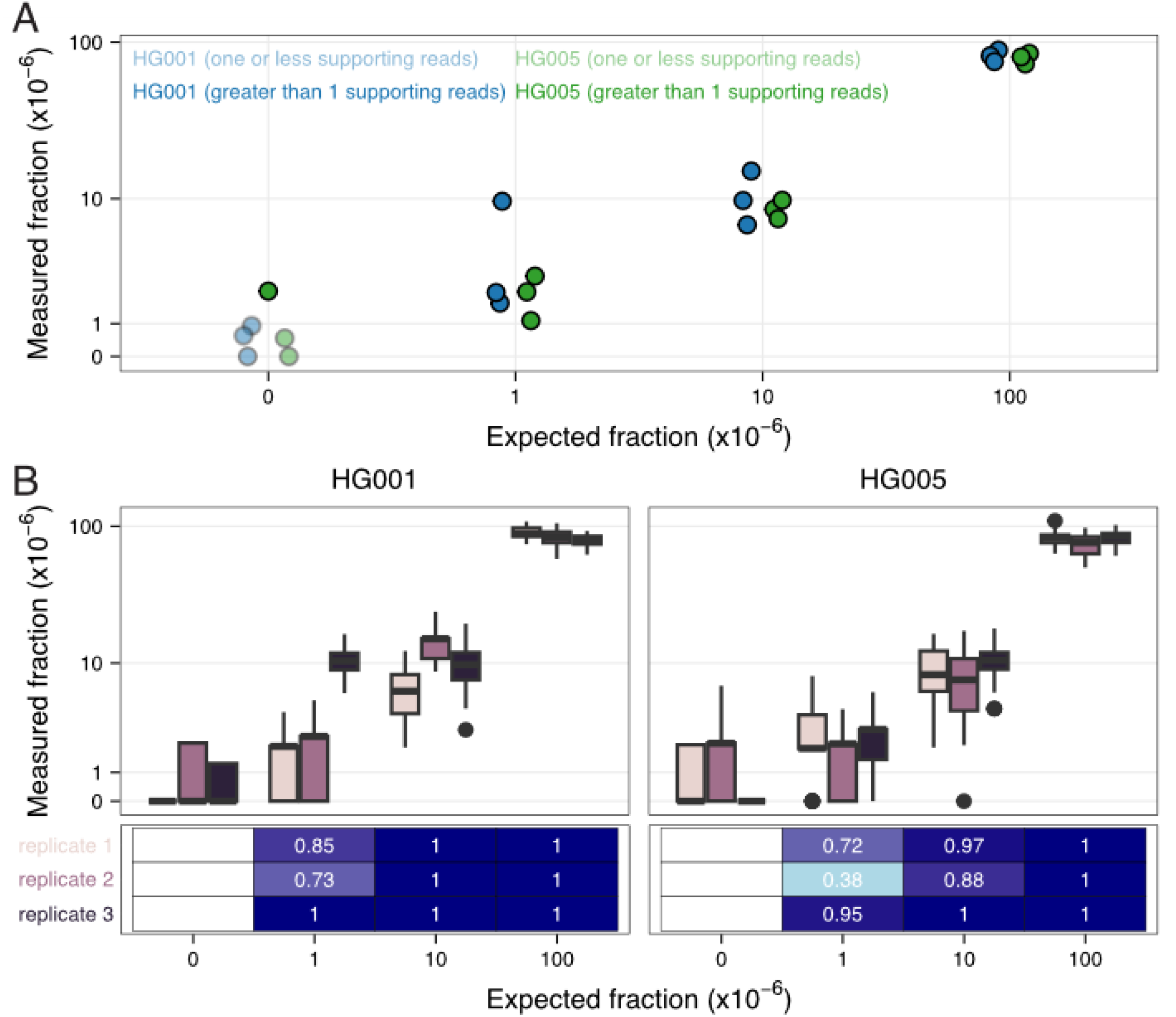
Benchmarking tumor fraction detection with ppmSeq using an in-vitro mixture of Genome in a Bottle (GIAB^*29*^) cell lines. **A** HG002A) 1 and HG005 were diluted into HG002 DNA in proportions of 1, 10 and 100 parts-per-million, and the fraction of reads supporting HG001- and HG005-unique-variants was measured in ppmSeq data across 3 repetitions. Measurement supported by reads are shown with high transparency. **B** Bootstrapping of the GIAB-unique signatures reveals high ppm-level accuracy. Each of the HG001- and HG005-unique homozygous variants were sampled a signature of size 30,000 SNVs, for 30 times with repeats. The distribution of GIAB-supporting reads is shown in boxplots, the area under ROC-curve, for each sample against the HG002 sample (expected fraction=0) is shown in heatmaps, separating between three experimental repetitions.

**Supplementary Figure 7.**
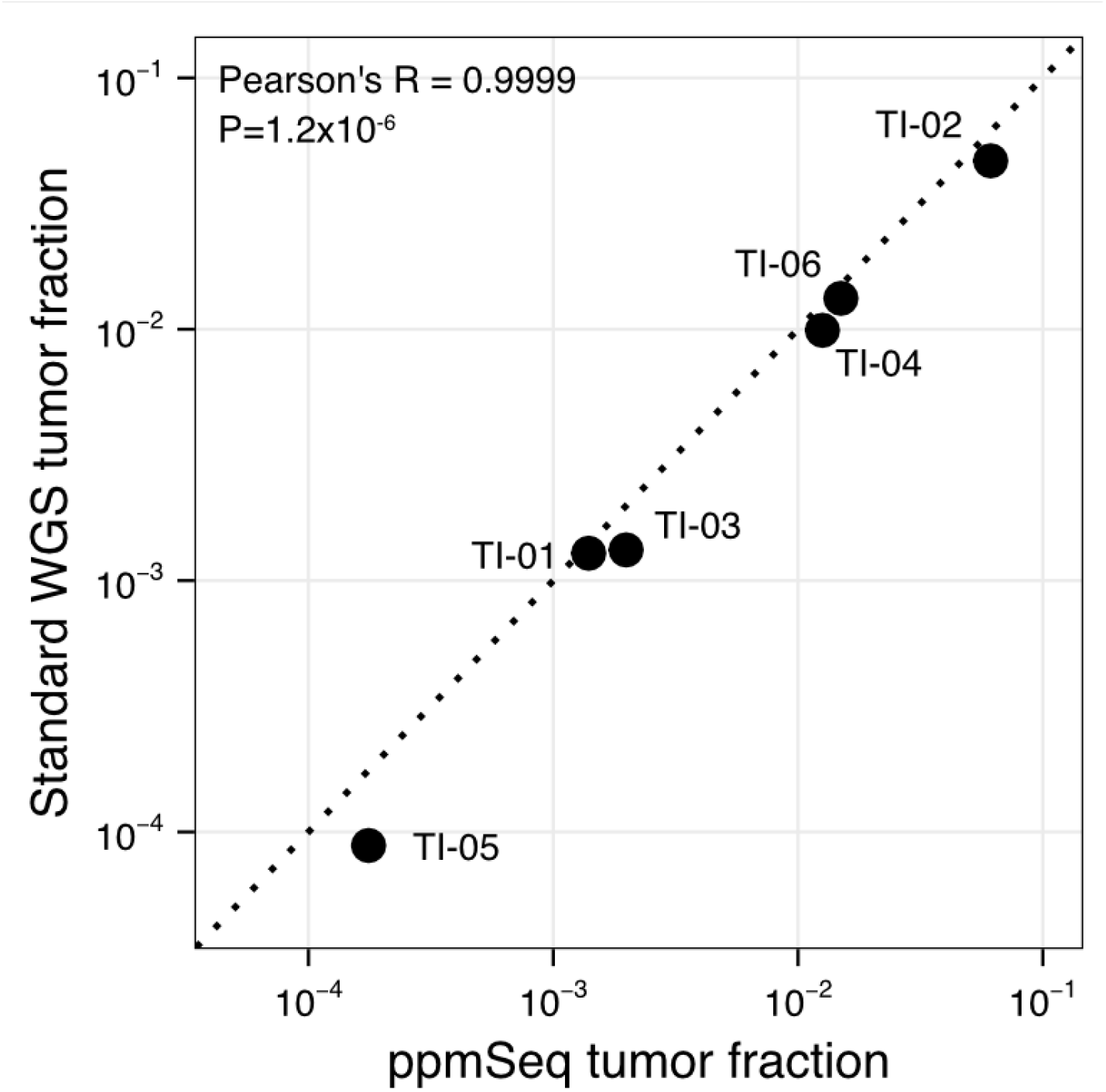
Tumor-informed tumor fractions in Ultima standard sequencing and ppmSeq datasets. Tumor informed tumor fractions measured from the same samples sequenced with both ppmSeq and standard Ultima Genomics sequencing. Patient number corresponding to details in **Supplementary Table 4** shown as text.

**Supplementary Figure 8.**
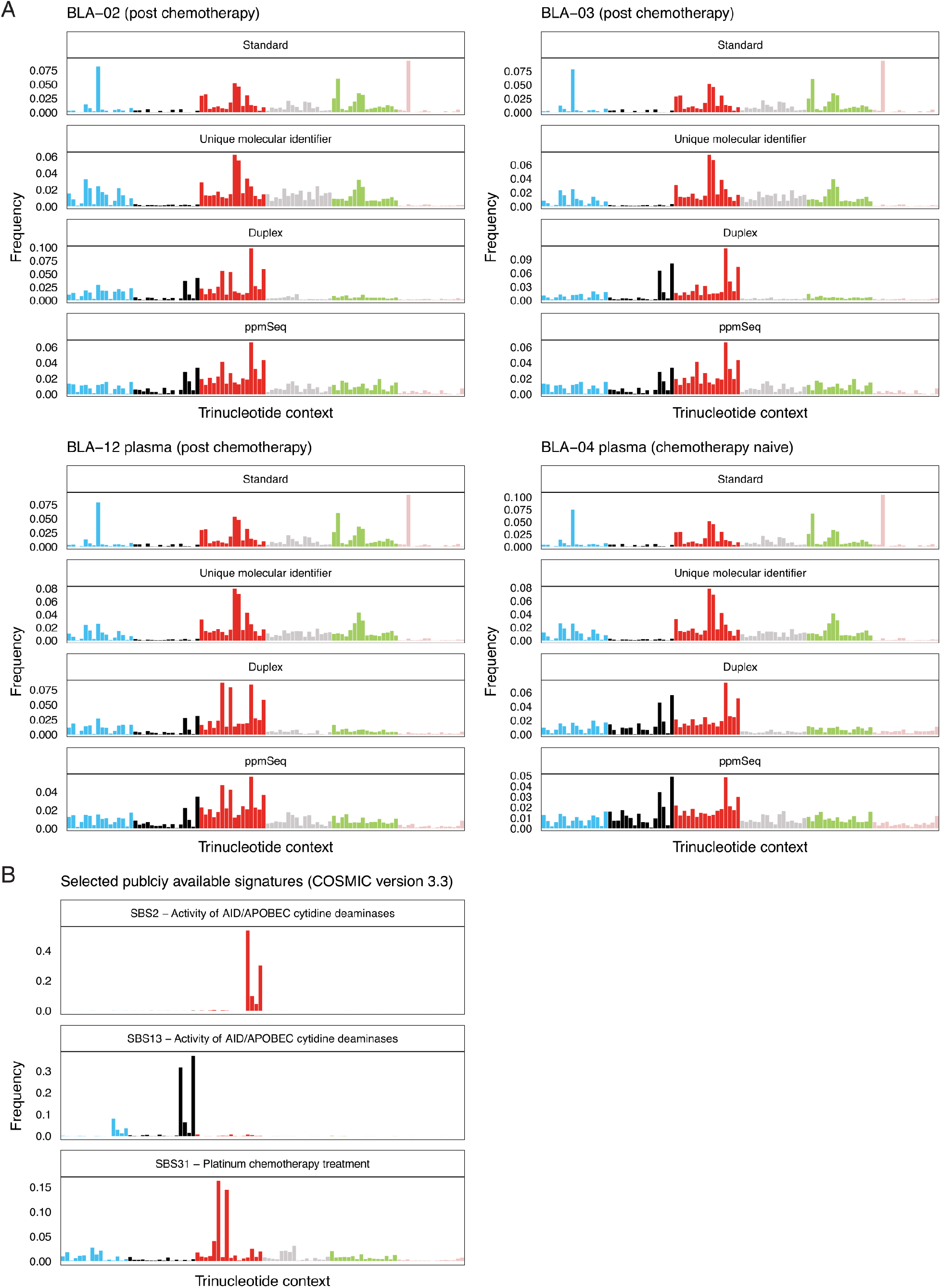
Trinucleotide frequencies patient plasma samples and previously described signatures. **A** Signatures in n = 4 high tumor burden cell-free DNA from patients with urothelial cancer. Tumor burden was considered high when ichorCNA tumor fraction estimates exceeded 5%. Primary signatures detected in Duplex and ppmSeq datasets were SBS2 and SBS13, and SBS31 in post-chemotherapy treated patients. **B** Previously described mutational signatures for SBS2, SBS13 and SBS31 (Cosmic v.3.3) are readily apparent in the patient plasma signatures.

## References

1. Coorens, T. H. H. et al. The somatic mutation landscape of normal gastric epithelium. Nature 640, 418–426 (2025).

2. Yokoyama, A. et al. Age-related remodelling of oesophageal epithelia by mutated cancer drivers. Nature 565, 312–317 (2019).

3. Olafsson, S. et al. Somatic Evolution in Non-neoplastic IBD-Affected Colon. Cell 182, 672–684.e11 (2020).

4. Brunner, S. F. et al. Somatic mutations and clonal dynamics in healthy and cirrhotic human liver. Nature 574, 538–542 (2019).

5. Zviran, A. et al. Genome-wide cell-free DNA mutational integration enables ultra-sensitive cancer monitoring. Nat Med 26, 1114–1124 (2020).

6. Cheng, A. P. et al. Error-corrected flow-based sequencing at whole-genome scale and its application to circulating cell-free DNA profiling. Nat Methods 1–9 (2025) doi:10.1038/s41592-025-02648-9.

7. Widman, A. J. et al. Ultrasensitive plasma-based monitoring of tumor burden using machine-learning-guided signal enrichment. Nat Med 30, 1655–1666 (2024).

8. Abascal, F. et al. Somatic mutation landscapes at single-molecule resolution. Nature 593, 405–410 (2021).

9. Hoang, M. L. et al. Genome-wide quantification of rare somatic mutations in normal human tissues using massively parallel sequencing. Proceedings of the National Academy of Sciences 113, 9846–9851 (2016).

10. Schmitt, M. W. et al. Detection of ultra-rare mutations by next-generation sequencing. Proceedings of the National Academy of Sciences 109, 14508–14513 (2012).

11. Bae, J. H. et al. Single duplex DNA sequencing with CODEC detects mutations with high sensitivity. Nat Genet 55, 871–879 (2023).

12. Almogy, G. et al. Cost-efficient whole genome-sequencing using novel mostly natural sequencing-by-synthesis chemistry and open fluidics platform. 2022.05.29.493900 Preprint at 10.1101/2022.05.29.493900 (2022).

13. Blauwkamp, T. A. et al. Analytical and clinical validation of a microbial cell-free DNA sequencing test for infectious disease. Nat Microbiol 4, 663–674 (2019).

14. Cheng, A. P. et al. A cell-free DNA metagenomic sequencing assay that integrates the host injury response to infection. Proceedings of the National Academy of Sciences 116, 18738–18744 (2019).

15. Burnham, P. et al. Urinary cell-free DNA is a versatile analyte for monitoring infections of the urinary tract. Nat Commun 9, 2412 (2018).

16. De Vlaminck, I. et al. Noninvasive monitoring of infection and rejection after lung transplantation. Proc Natl Acad Sci U S A 112, 13336–13341 (2015).

17. Chiu, R. W. K. et al. Noninvasive prenatal diagnosis of fetal chromosomal aneuploidy by massively parallel genomic sequencing of DNA in maternal plasma. Proc Natl Acad Sci U S A 105, 20458–20463 (2008).

18. Fan, H. C., Blumenfeld, Y. J., Chitkara, U., Hudgins, L. & Quake, S. R. Noninvasive diagnosis of fetal aneuploidy by shotgun sequencing DNA from maternal blood. Proceedings of the National Academy of Sciences 105, 16266–16271 (2008).

19. Cohen, J. D. et al. Detection of low-frequency DNA variants by targeted sequencing of the Watson and Crick strands. Nat Biotechnol 39, 1220–1227 (2021).

20. Phallen, J. et al. Direct detection of early-stage cancers using circulating tumor DNA. Science Translational Medicine 9, eaan2415 (2017).

21. Chaudhuri, A. A. et al. Early Detection of Molecular Residual Disease in Localized Lung Cancer by Circulating Tumor DNA Profiling. Cancer Discovery 7, 1394–1403 (2017).

22. Black, J. R. M. et al. Ultrasensitive ctDNA detection for preoperative disease stratification in early-stage lung adenocarcinoma. Nat Med 31, 70–76 (2025).

23. Newman, A. M. et al. An ultrasensitive method for quantitating circulating tumor DNA with broad patient coverage. Nat Med 20, 548–554 (2014).

24. Newman, A. M. et al. Integrated digital error suppression for improved detection of circulating tumor DNA. Nat Biotechnol 34, 547–555 (2016).

25. Kurtz, D. M. et al. Enhanced detection of minimal residual disease by targeted sequencing of phased variants in circulating tumor DNA. Nat Biotechnol 39, 1537–1547 (2021).

26. Meddeb, R. et al. Quantifying circulating cell-free DNA in humans. Sci Rep 9, 5220 (2019).

27. Liu, M. H. et al. DNA mismatch and damage patterns revealed by single-molecule sequencing. Nature 630, 752–761 (2024).

28. Jiang, P. et al. Detection and characterization of jagged ends of double-stranded DNA in plasma. Genome Res. https://doi.org/10.1101/gr.261396.120 (2020) doi:10.1101/gr.261396.120.

29. Genome in a bottle—a human DNA standard. Nature Biotechnology 33, 675–675 (2015).

30. Aaltonen, L. A. et al. Pan-cancer analysis of whole genomes. Nature 578, 82–93 (2020).

31. Sethi, H. et al. Abstract 4542: Analytical validation of the SignateraTM RUO assay, a highly sensitive patient-specific multiplex PCR NGS-based noninvasive cancer recurrence detection and therapy monitoring assay. Cancer Res 78, 4542 (2018).

32. Campbell, P. J. et al. Pan-cancer analysis of whole genomes. Nature 578, 82–93 (2020).

33. Moding, E. J., Nabet, B. Y., Alizadeh, A. A. & Diehn, M. Detecting liquid remnants of solid tumors: circulating tumor DNA minimal residual disease. Cancer Discov 11, 2968–2986 (2021).

34. Black, J. R. M. et al. Longitudinal ultrasensitive ctDNA monitoring for high-resolution lung cancer risk prediction. Cell 188, 7083–7098.e18 (2025).

35. Tie, J. et al. Circulating Tumor DNA Analysis Guiding Adjuvant Therapy in Stage II Colon Cancer. N Engl J Med 386, 2261–2272 (2022).

36. Shaw, J. A. et al. Serial Postoperative Circulating Tumor DNA Assessment Has Strong Prognostic Value During Long-Term Follow-Up in Patients With Breast Cancer. JCO Precis Oncol 8, e2300456 (2024).

37. Cindy Yang, S. Y. et al. Pan-cancer analysis of longitudinal metastatic tumors reveals genomic alterations and immune landscape dynamics associated with pembrolizumab sensitivity. Nat Commun 12, 5137 (2021).

38. Adalsteinsson, V. A. et al. Scalable whole-exome sequencing of cell-free DNA reveals high concordance with metastatic tumors. Nat Commun 8, 1324 (2017).

39. Nguyen, D. D. et al. The Interplay between Mutagenesis and Extrachromosomal DNA Shapes Urothelial Cancer Evolution. 2023.05.07.538753 Preprint at 10.1101/2023.05.07.538753 (2023).

40. Petljak, M. et al. Characterizing Mutational Signatures in Human Cancer Cell Lines Reveals Episodic APOBEC Mutagenesis. Cell 176, 1282–1294.e20 (2019).

41. Osorio, F. G. et al. Somatic Mutations Reveal Lineage Relationships and Age-Related Mutagenesis in Human Hematopoiesis. Cell Reports 25, 2308–2316.e4 (2018).

42. Alexandrov, L. B. et al. Signatures of mutational processes in human cancer. Nature 500, 415–421 (2013).

43. Moore, L. et al. The mutational landscape of human somatic and germline cells. Nature 597, 381–386 (2021).

44. Cho, S. W. et al. Analysis of off-target effects of CRISPR/Cas-derived RNA-guided endonucleases and nickases. Genome Res 24, 132–141 (2014).

45. Schmitt, M. W., Loeb, L. A. & Salk, J. J. The influence of subclonal resistance mutations on targeted cancer therapy. Nat Rev Clin Oncol 13, 335–347 (2016).

46. Rossi, D. et al. Clinical impact of small TP53 mutated subclones in chronic lymphocytic leukemia. Blood 123, 2139–2147 (2014).

47. Landau, D. A. et al. Evolution and impact of subclonal mutations in chronic lymphocytic leukemia. Cell 152, 714–726 (2013).

48. Herberts, C. et al. Deep whole-genome ctDNA chronology of treatment-resistant prostate cancer. Nature 608, 199–208 (2022).

49. Wan, J. C. M. et al. ctDNA monitoring using patient-specific sequencing and integration of variant reads. Science Translational Medicine 12, eaaz8084 (2020).

50. Murtaza, M. et al. Non-invasive analysis of acquired resistance to cancer therapy by sequencing of plasma DNA. Nature 497, 108–112 (2013).

51. Cristiano, S. et al. Genome-wide cell-free DNA fragmentation in patients with cancer. Nature 570, 385–389 (2019).

52. Bettegowda, C. et al. Detection of Circulating Tumor DNA in Early-and Late-Stage Human Malignancies. Sci Transl Med 6, 224ra24 (2014).

53. Tie, J. et al. Circulating Tumor DNA Analysis Guiding Adjuvant Therapy in Stage II Colon Cancer. New England Journal of Medicine 386, 2261–2272 (2022).

54. Shao, D. D. et al. Advances in single-cell DNA sequencing enable insights into human somatic mosaicism. Nat Rev Genet https://doi.org/10.1038/s41576-025-00832-3 (2025) doi:10.1038/s41576-025-00832-3.

55. Martincorena, I. et al. High burden and pervasive positive selection of somatic mutations in normal human skin. Science 348, 880–886 (2015).

56. Martincorena, I. et al. Universal Patterns of Selection in Cancer and Somatic Tissues. Cell 171, 1029–1041.e21 (2017).

57. Martincorena, I. & Campbell, P. J. Somatic mutation in cancer and normal cells. Science 349, 1483–1489 (2015).

58. Pel, J. et al. Duplex Proximity Sequencing (Pro-Seq): A method to improve DNA sequencing accuracy without the cost of molecular barcoding redundancy. PLOS ONE 13, e0204265 (2018).

59. Lawson, A. R. J. et al. Somatic mutation and selection at epidemiological scale. 2024.10.30.24316422 Preprint at 10.1101/2024.10.30.24316422 (2024).

60. Tie, J. et al. Circulating Tumor DNA Analysis Guiding Adjuvant Therapy in Stage II Colon Cancer. N Engl J Med 386, 2261–2272 (2022).

61. Powles, T. et al. ctDNA guiding adjuvant immunotherapy in urothelial carcinoma. Nature 595, 432–437 (2021).

62. Bratman, S. V. et al. Personalized circulating tumor DNA analysis as a predictive biomarker in solid tumor patients treated with pembrolizumab. Nat Cancer 1, 873–881 (2020).

63. Coorens, T. H. H. et al. The somatic mutation landscape of normal gastric epithelium. Nature 640, 418–426 (2025).

64. Lawson, A. R. J. et al. Extensive heterogeneity in somatic mutation and selection in the human bladder. Science 370, 75–82 (2020).

65. Moore, L. et al. The mutational landscape of normal human endometrial epithelium. Nature 580, 640–646 (2020).

66. Anglesio, M. S. et al. Cancer-Associated Mutations in Endometriosis without Cancer. N Engl J Med 376, 1835–1848 (2017).

67. Lee-Six, H. et al. The landscape of somatic mutation in normal colorectal epithelial cells. Nature 574, 532–537 (2019).

68. Martincorena, I. et al. Somatic mutant clones colonize the human esophagus with age. Science 362, 911–917 (2018).

69. Mitchell, E. et al. Clonal dynamics of haematopoiesis across the human lifespan. Nature 606, 343–350 (2022).

70. Jaiswal, S. et al. Age-related clonal hematopoiesis associated with adverse outcomes. N Engl J Med 371, 2488–2498 (2014).

71. Ultimagen. healthomics-workflows/docs/UG_ cram_format.pdf at main · Ultimagen/healthomics-workflows. GitHub https://github.com/Ultimagen/healthomics-workflows/blob/main/docs/UG_cram_format.pdf.

72. Chen, T. & Guestrin, C. XGBoost: A Scalable Tree Boosting System. in Proceedings of the 22nd ACM SIGKDD International Conference on Knowledge Discovery and Data Mining 785–794 (Association for Computing Machinery, New York, NY, USA, 2016). doi:10.1145/2939672.2939785.

73. Phan, L. et al. The evolution of dbSNP: 25 years of impact in genomic research. Nucleic Acids Research 53, D925–D931 (2025).

74. Chen, S. et al. A genomic mutational constraint map using variation in 76,156 human genomes. Nature 625, 92–100 (2024).

75. Lundberg, S. M. & Lee, S.-I. A Unified Approach to Interpreting Model Predictions. in Advances in Neural Information Processing Systems vol. 30 (Curran Associates, Inc., 2017).

76. Zook, J. M. et al. Extensive sequencing of seven human genomes to characterize benchmark reference materials. Sci Data 3, 160025 (2016).

77. Jin, H. et al. Accurate and sensitive mutational signature analysis with MuSiCal. Nat Genet 56, 541–552 (2024).

78. Sanchez, C. et al. Circulating nuclear DNA structural features, origins, and complete size profile revealed by fragmentomics. JCI Insight 6, e144561.

